# Mechanisms of maternal antibody interference to rotavirus vaccination

**DOI:** 10.1101/2025.02.26.640336

**Authors:** Tawny L Chandler, Sarah Woodyear, Valerie Chen, Tom Lonergan, Natalie Baker, Katherine Harcourt, Simon Clare, Faraz Ahmed, Sarah L Caddy

**Affiliations:** Baker Institute for Animal Health, Cornell University, Ithaca, NY, United States; Department of Medicine, University of Cambridge, Cambridge, UK; Genomics Innovation Hub and TREx Facility, Institute of Biotechnology, Cornell University, Ithaca, NY, United States; Department of Microbiology and Immunology, College of Veterinary Medicine, Cornell University, Ithaca, NY, United States

**Author notes:** These authors contributed equally (Tawny L Chandler, Sarah Woodyear).

## Abstract

Maternal antibodies (MatAbs) are transferred transplacentally during pregnancy and through breast milk after birth to provide protection whilst the neonatal immune response is immature. However, MatAbs also suppress the development of neonatal B cell responses via mechanisms that are not well defined. MatAbs can therefore result in poor vaccine performance in the infant, placing them at risk against potentially life-threatening pathogens such as rotavirus. It is essential we understand the mechanisms by which MatAbs interact with neonatal immunity, so that strategies can be developed to overcome this major vaccine issue.

To investigate mechanisms of interference we developed a mouse model of neonatal oral rotavirus vaccination in the presence and absence of MatAbs. Oral vaccination with attenuated murine rotavirus induced robust neonatal antibody responses, whereas vaccination failed to induce seroconversion in the presence of MatAbs. Vaccination with heterologous strains, reduced MatAb titers and vaccination in FcγRIIB knockout mice did not overcome interference. However, live vaccine replication was blocked in the presence of MatAbs and more rapid waning of MatAbs was observed following vaccination. This is indicative of premature vaccine clearance and likely reduced antigen encounter by B cells. Single-cell RNA sequencing of mesenteric lymph nodes revealed diminished plasma and germinal center B cell subpopulations as well as global reduction of interferon-stimulated genes in the presence of MatAbs. Our model has also enabled identification of strategies to reduce the effects of interference.

In summary, we have tested multiple hypotheses for MatAb-mediated interference to rotavirus vaccination in a mouse model, and demonstrated that premature FcγRIIB signaling and epitope masking are not the primary mechanisms of vaccine failure. Rather our data supports the conclusion that MatAb-mediated vaccine clearance is a key mechanism of interference to oral rotavirus vaccine.

## Introduction

Neonates of all mammalian species are considered immunologically naïve at birth. This places the neonate at a high risk of infection whilst their immune system is maturing. To counter this, mothers have evolved a strategy to deliver immune protection to their offspring in the form of maternal antibodies (MatAbs). Antibodies in the maternal circulation can be passively transferred to the fetus across the placenta, and postnatally to the neonate in milk. These MatAbs function to protect the naïve offspring against pathogens faced in the environment. Lower levels of MatAbs can be correlated with increased infectious disease in the infant (1).

Whilst MatAbs are an undoubted evolutionary advantage, a paradoxical effect of their presence is a reduction in vaccine efficacy in infants. Often known as MatAb interference or MatAb blunting, the phenomenon of lower vaccine efficacy in the face of MatAbs has been described for many different vaccines. This includes numerous viruses of human and veterinary importance, including measles virus (2), foot and mouth disease virus (3), hepatitis A virus (4), and poliovirus (5). MatAb interference is especially concerning for rotavirus vaccines. Rotavirus is a significant cause of gastroenteritis in young children, attributed to over 200,000 deaths in children under five years old (6). The current rotavirus vaccines are live attenuated strains that were first licensed in 2006, and although these vaccines have proven to be highly effective in high-income countries, their efficacy is often less than 50% in low- and middle-income regions. Multiple factors have been proposed to be responsible for this poor efficacy, including malnutrition, co-infections and host genetics, but a leading explanation is MatAb interference (7–9). MatAb titers are generally higher in lower-middle-income countries which could account for this geographical variation in vaccine efficacy (10–12). A substantial number of clinical studies have identified a negative association between high titers of MatAbs and poor seroconversion following vaccination with currently used rotavirus vaccines (13–18).

The mechanisms by which MatAbs can reduce vaccine efficacy have been of significant interest over many years yet remain unclear. A number of different theories have been proposed, including MatAb-mediated masking of vaccine epitopes, and direct inhibition of neonatal B cells via MatAbs cross linking the B cell receptor (BCR) and the inhibitory FcγRIIB receptor (1, 19). Clearance of vaccines by MatAbs so the neonatal immune response remains naïve has also been a leading theory, but some studies have shown that B cell activation still occurs in the presence of high levels of MatAbs (20). Ultimately it is challenging to apply conclusions drawn from one experimental vaccine model to another, especially given the diversity in immune responses required to induce effective protection to different pathogens and different types of vaccine.

In this study, we aimed to understand the role of MatAbs in the context of rotavirus vaccination, given the widely reported issues with rotavirus vaccine efficacy in human infants. Prior research in the field of MatAb interference mechanisms has largely focused on systemic or respiratory pathogens (20, 21), and we predicted that responses to a gastrointestinal pathogen and orally delivered vaccine would be distinct. Here, we successfully established a mouse model of MatAb interference to a live attenuated rotavirus vaccine, and used this to investigate the potential mechanisms responsible for poor vaccine efficacy in the presence of MatAbs. Using a combination of hypothesis-driven experimental approaches we have shown that the major mechanism of interference is MatAb-driven clearance of the rotavirus vaccine prior to vaccine encounter by the neonatal immune system.

## Results

### Rotavirus-specific maternal antibodies block seroconversion to rotavirus vaccination in pups

Rotavirus-specific MatAbs in humans have been associated with reduced infant seroconversion to oral rotavirus vaccines in multiple clinical studies (13–18). To study the immunological mechanisms underpinning this observation we aimed to establish a mouse model of rotavirus-specific MatAb transfer and vaccination. We began by infecting adult female C57BL/6 mice with 10 FFU of an attenuated murine rotavirus strain EMcN known to readily infect this mouse line (22). Infection was performed by oral gavage, with control mice receiving an equal volume of PBS. Female mice were mated with male BALB/c mice after seroconversion was confirmed by ELISA at 7-10 days post infection. C57BL/6 x BALB/c pups were vaccinated at 7 days old using a subclinical dose of EMcN rotavirus delivered by oral gavage. This was designed to model administration of live attenuated oral vaccines to human infants at 8 weeks old. All pups within a litter received the same treatment and both genders were considered in analyses. Pups were weaned at 3 weeks old, and blood samples were collected every 2 weeks from 4-10 weeks of age. The experimental pipeline is presented in Figure 1A.

**Figure 1.**
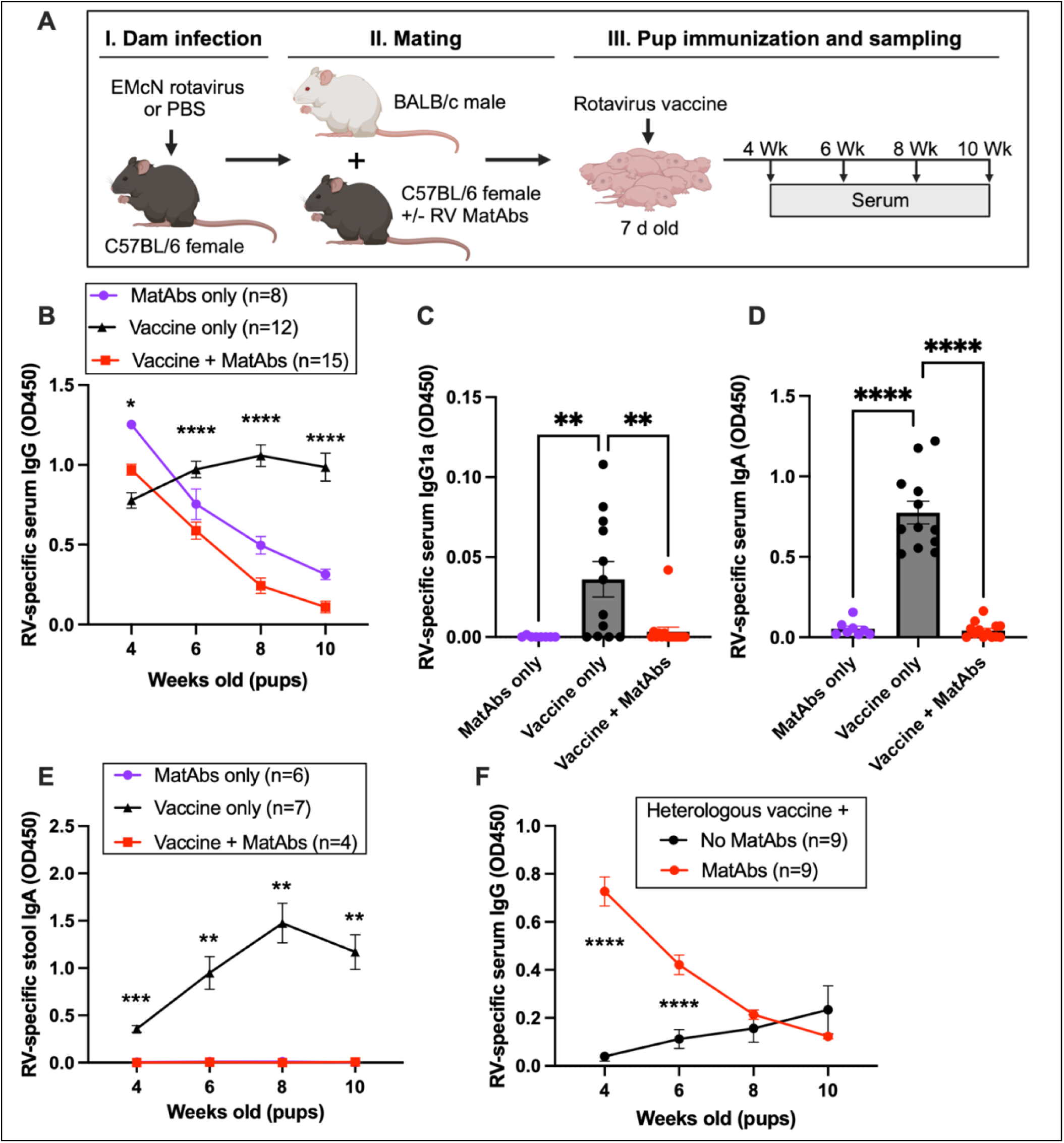
Maternal antibodies interfere with pup immune responses. (**A**) Schematic diagram of experimental protocol. (**B**) Serum rotavirus-specific IgG in pups as measured by ELISA from 4-10 weeks of age. Differentiation of maternal and pup antibodies by (**C**) IgG1a and (**D**) IgA at 10 weeks of age. (**E**) Stool IgA titers in the presence or absence of MatAbs. (**F**) Serum IgG in pups following vaccination with a heterologous rotavirus vaccine strain. Statistical significance was determined by two-way ANOVA with repeated measures (**B**, **E** and **F**) or one-way ANOVA (**C** and **D**). Significant Tukey’s adjusted pair-wise comparisons of vaccination in the absence or presence of MatAbs (**B** and **E**) and Bonferonni-corrected pair-wise comparisons (**C**, **D** and **F**) are shown (* *p* ≤ 0.05; ** *p* < 0.01, *** *p* < 0.001, **** *p* < 0.0001).

Serum antibodies circulating in the pups post-weaning were quantified by a rotavirus-specific sandwich ELISA. As shown in Figure 1B, rotavirus-specific IgG from the dam was readily detectable in a litter of pups born to a seropositive dam (‘MatAbs only’) at 4 weeks of age. These MatAbs were then observed to wane over the subsequent 6-week period. A second litter of pups that were vaccinated with live attenuated murine rotavirus in the absence of any MatAbs (‘Vaccine only’) made a robust rotavirus- specific IgG response. However, a third litter of pups that received murine rotavirus vaccine in the presence of MatAbs (‘Vaccine + MatAbs’) failed to mount their own detectable serum IgG response. This showed that in our experimental model MatAbs negatively affected the ability of pups to seroconvert post- vaccination, in agreement with observations made in human studies.

One advantage of using a mouse model over human samples is the ability to differentiate MatAbs detected in pup serum from those made by the pup themselves. We used two alternative methods to verify that pups vaccinated in the presence of MatAbs did not produce a detectable antibody response. Firstly, our use of C57BL/6 dams and BALB/c males took advantage of variation in IgG1 alleles within these two mouse strains. BALB/c mice have the IgG1a allele, whereas C57BL/6 mice have IgG1b. Use of an IgG1a-specific secondary antibody to quantify serum antibodies by ELISA only detected IgG1 produced by the pup. As shown in Figure 1C, when pups were 10 weeks old, it was clear that only pups receiving the oral vaccine in the absence of MatAbs mounted their own IgG1 response. As rotavirus is a mucosal pathogen, an alternative means of differentiating maternal and neonatal antibody responses is by quantification of IgA. IgG from the mother’s milk can enter the pup circulation due to FcRn-mediated uptake across the intestinal barrier (23), but IgA does not cross the intestinal epithelium, thus any IgA detected in pup serum must be produced by the pup themselves. Figure 1D shows serum IgA of 10- week-old mice born to mothers with or without antibodies, and mirrors Figure 1C accordingly. We also wanted to verify that production of IgA detected in stool was similarly impacted by the presence of MatAbs. To achieve this we independently vaccinated a second cohort of litters with and without MatAbs, and longitudinally collected stool samples. IgA ELISA results presented in Figure 1E again show no stool IgA is detected in the presence of MatAbs. Overall, these results support the conclusion that MatAbs interfere with the ability of the pup to seroconvert to rotavirus vaccination.

Many strains of human rotavirus circulate in human populations, yet rotavirus vaccines are currently restricted to either a single strain (G1P[8] e.g. Rotarix, GSK), or five reassortant strains (e.g. Rotateq, Merck). This means that MatAbs transferred are likely to target rotavirus strains that differ from those in the vaccine administered to the neonate. To address whether MatAb interference can still occur in our model when the vaccinating strain and MatAb specificity are mismatched, we tested a heterologous vaccine approach. Dams used were seropositive to the murine rotavirus strain (G16P[16]) as before, whereas pups were vaccinated with a G1P[8] human rotavirus strain. We have previously shown that murine rotavirus-specific antibodies can neutralize human rotavirus strains, albeit to a lesser degree than for homologous strains (24). As shown in Figure 1F, no pups showed evidence of seroconversion to the human rotavirus vaccine in the presence of MatAbs, whereas some degree of seroconversion was apparent in the naïve pups. It is important to note that the limited seroconversion identified in the no MatAb group is likely due to the species-specificity of rotaviruses (24, 25). Further experiments in this study therefore continued to focus on murine-specific rotaviruses to maximise potential seroconversion.

### Germinal center formation is limited in the presence of maternal antibodies

Whilst no pup antibody was detected in the serum of pups vaccinated in the presence of MatAbs, we questioned what effect MatAbs were having on the wider neonatal B cell response. To address this, we repeated our experimental timeline with administration of rotavirus or PBS to dams, transfer of any induced MatAbs to pups, and then vaccination of pups at 7 days of age. Pups were culled 20 days post vaccination and mesenteric lymph nodes (MLN) were collected from all mice for analysis by flow cytometry or histology.

We observed a significant decrease in the formation of germinal centers in MLN of mice vaccinated in the presence of MatAbs. This was apparent via flow cytometry where a statistically significant difference was detected (Figure 2A, germinal centers identified by B220, GL7 and Fas staining). We verified this result using both haematoxylin and eosin (H&E) staining (Figure 2B) and immunofluorescence (staining for Ki67 and B220, Figure 2C) of MLN tissue sections from pups in the same litters. Minimal visible germinal center formation was evident in pups vaccinated in the presence of MatAbs. This is in agreement with the observed serological responses, but in contrast to findings with a MatAb model and parenteral influenza subunit vaccination (20). Interestingly there was no difference in the relative level of T follicular helper (TfH) cells in MLNs (Figure 2D). Gating strategies for GC and TfH cells are outlined in Supplementary Figure 1.

**Figure 2.**
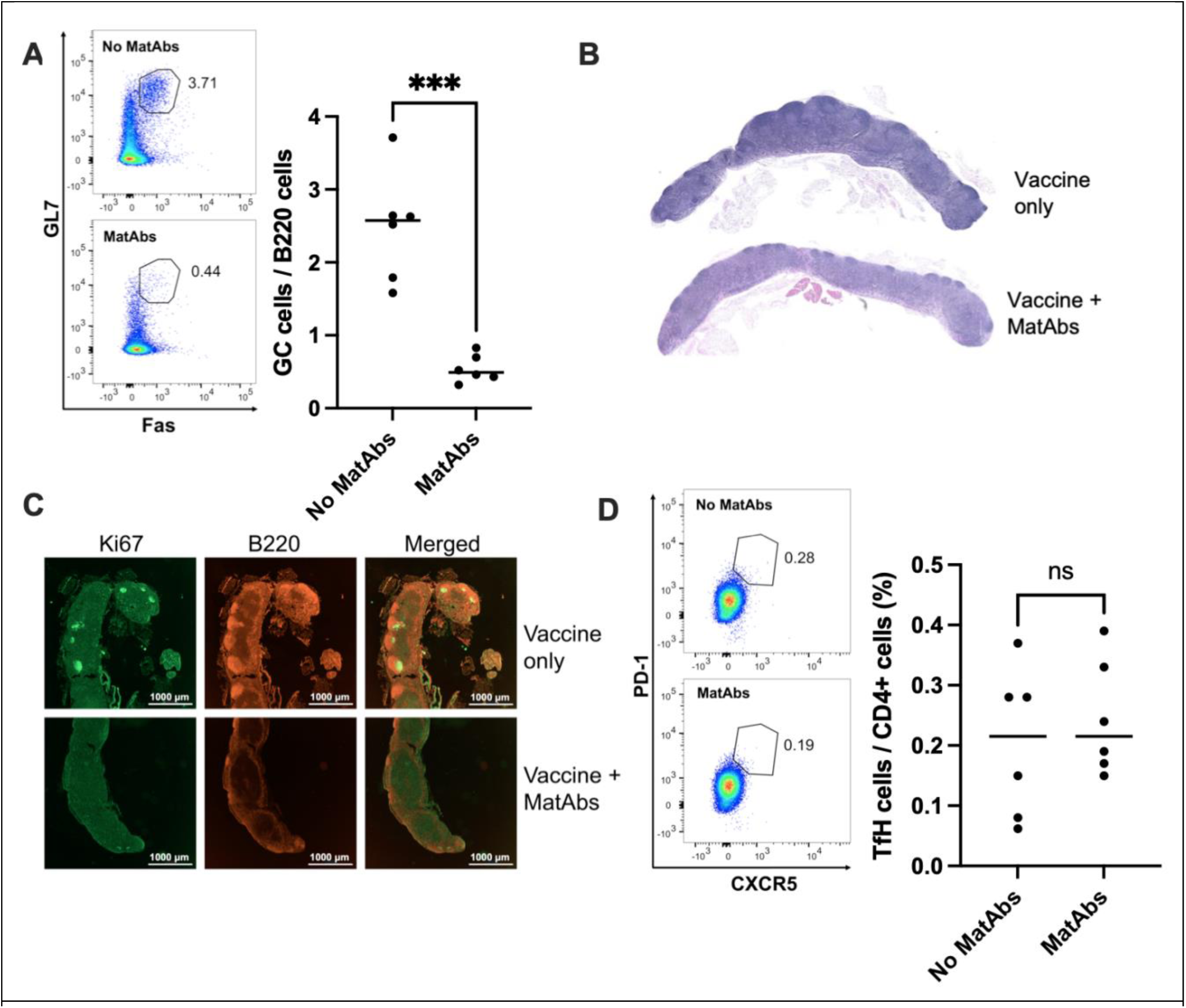
Maternal antibodies limit formation of germinal centers (GCs) in draining lymph nodes. (**A**) Flow cytometry of GCs in mesenteric lymph nodes (MLNs) from pups 20 days post vaccination, with representative flow cytometry plots and scatterplot presenting all mice. (**B**) H&E staining and (**C**) immunofluorescence of MLN sections from mice vaccinated in the presence/absence of MatAbs. (**D**) Flow cytometry of T follicular helper cells in MLNs. Statistical significance was determined by unpaired two-tailed t-tests (*** *p* < 0.001).

### Maternal antibody-mediated interference is robust with very low maternal antibody doses

Our first series of experiments demonstrated complete ablation of pup seroconversion to rotavirus vaccination in the presence of MatAbs induced by infection of dams with high doses of rotavirus. As MatAb titer has been correlated with seroconversion to rotavirus vaccination in human infants (13–17), we next asked whether pup seroconversion could be achieved by lowering the dose of MatAbs received by pups. We predicted that reducing the titer of MatAbs in the dam would enable more pups to seroconvert. This would facilitate the comparison of antibody repertoires between the dam and the pups to examine the proposed phenomenon of MatAb-mediated epitope masking.

To lower the dose of MatAbs received by the pups we used two alternative approaches. Firstly, we infected the dams with two different doses of rotavirus one week prior to mating, with the expectation that maternal rotavirus-specific antibody titers would be different during pregnancy and lactation. Figure 3A presents maternal serum IgG and IgA quantified by ELISA in two dams infected with rotavirus, showing how circulating MatAbs changed following the birth of pups. Whilst serum IgA was clearly distinct between the two dams, inducing different levels of serum IgG was more challenging, and vaccination of pups appears to have a boosting effect on serum antibody levels in the dam. Analysis of serum rotavirus - specific IgG and IgA levels in pups born to the dams depicted in Figure 3A are shown in Figure 3B and 3C. We found that regardless of dam serum antibody levels, neither litter of pups with MatAbs seroconverted following rotavirus vaccination. However, as we observed that maternal IgG titers converged and likely reached a ceiling, separating MatAb delivery titer by this approach was shown to be an imperfect strategy.

**Figure 3.**
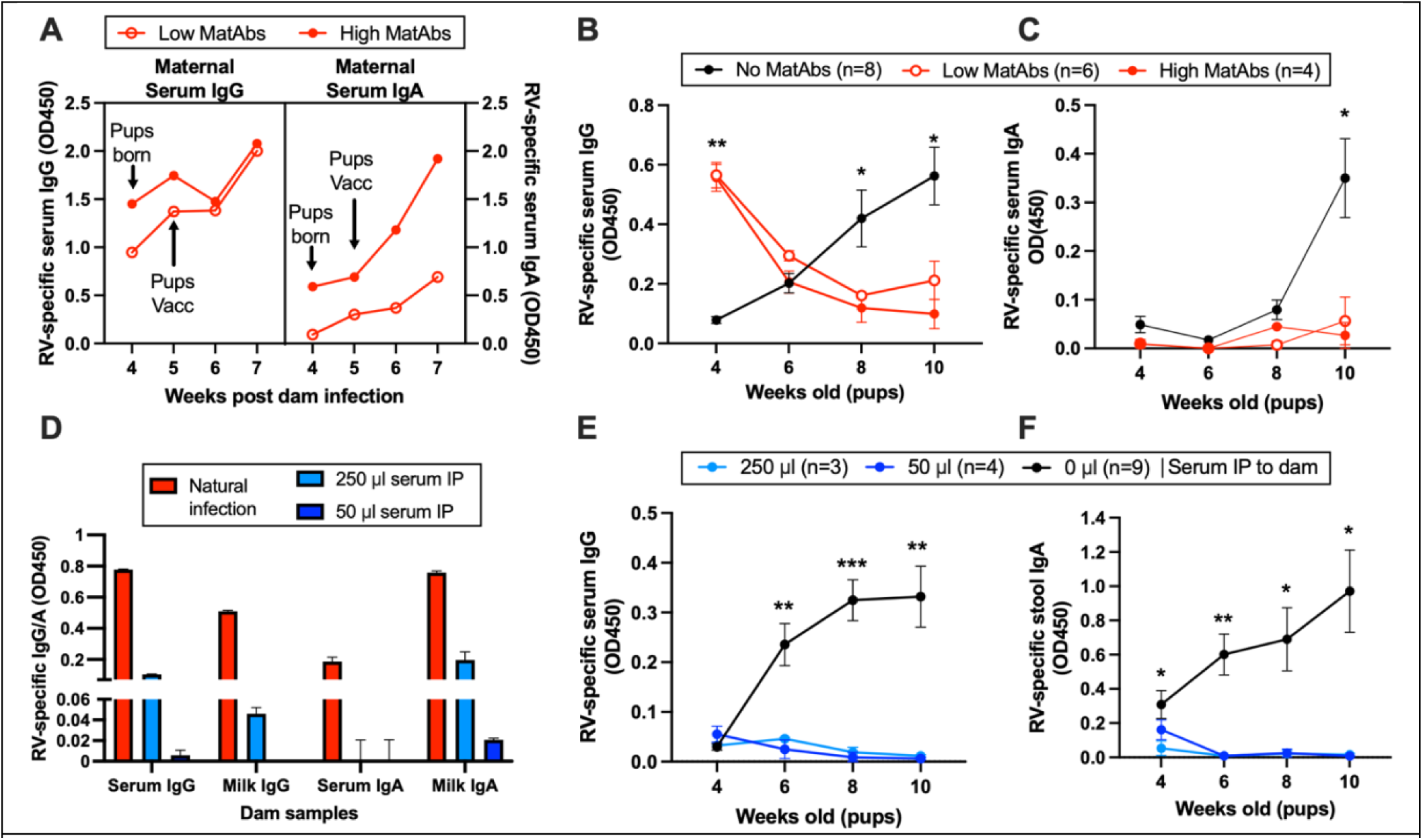
Maternal antibody-mediated interference is robust with low antibody doses. (**A**) Longitudinal measurement of dam rotavirus-specific antibody by ELISA. Longitudinal measurement of serum rotavirus-specific IgG (**B**) and IgA (**C**) in pups receiving high, low or no MatAbs. (**D**) Quantification of rotavirus-specific IgG and IgA at the time of pup vaccination in the serum and milk of dams naturally infected or receiving pooled serum from infected mice intraperitoneally (IP). Longitudinal measurement of rotavirus-specific serum IgG (**E**) and stool IgA (**F**) in pups receiving high, low or no MatAbs. Error bars show standard error of the mean, representing biological replicates (**B**, **C**, **D** and **E**) or technical replicates of a single mouse (**A** and **D**). Statistical significance was determined by two-way ANOVA with repeated measures (**B**, **C**, **E** and **F**). The largest significant Tukey-adjusted pair-wise comparisons between the absence or presence of MatAbs (**B** and **C**) and the absence of MatAb or passively transferred serum are shown (**E** and **F**) (* *p* ≤ 0.05; ** *p* < 0.01, *** *p* < 0.001, **** *p* < 0.0001).

For our second approach we used a passive transfer model, whereby a controlled amount of MatAb was delivered to dams. This allowed us to titrate down how much MatAb was transferred to pups. To achieve this, serum from rotavirus-specific antibody positive mice was pooled, and administered intraperitoneally (IP) to dams 24 hours prior to vaccination of pups at 7 days of age and a second time one-week post-vaccination. A key difference using this experimental approach as opposed to naturally induced MatAbs was that MatAb delivery via passive transfer was only via milk. Verification of rotavirus-specific antibody transfer from the intraperitoneal space to the maternal circulation and milk is shown in Figure 3D. Antibody titers in naturally infected dams is included for comparison. A log10 scale was used for the x-axis to demonstrate that IgA was detectable in milk of the dam receiving 50 μl serum albeit at a very low level. Unexpectedly, as shown in Figure 3E, no seroconversion was detectable in either litter of pups receiving MatAbs from passively transferred dams. As rotavirus-specific serum IgA responses were low in all pups (Figure S2A), rotavirus-specific IgA was also quantified in stool pellets collected from individual mice at each time point. Again, no seroconversion was demonstrated if pooled serum was administered to dams (Figure 3F). Overall, this shows that even very low titers of MatAbs delivered only by milk can interfere with infant vaccine responses in our model.

### FcγRIIB signaling is not a major mediator of maternal antibody interference

One of the major mechanisms proposed for MatAb interference has been premature signaling through inhibitory FcγRIIB (1, 19, 26). It has been theorized that MatAbs can function to co-ligate both the BCR and FcγRIIB on infant B cells simultaneously, resulting in blockade of B cell activation. To test this hypothesis, we replicated our mouse model of maternal interference to rotavirus vaccination in FcγRIIB knockout (KO) mice (kindly gifted from Jeffrey Ravetch, Rockefeller). FcγRIIB KO mice have been reported to have higher antibody titers due to the absence of signaling to bring an initial B cell response to a close (27), we therefore sought to evaluate any potential differences in rotavirus-specific IgG titers following primary infection. As shown in Figure 4A, both wild-type (WT) and KO mice exhibited a steady increase in IgG titers over the 5-week period post infection. A comparable titer of MatAb was therefore expected to be delivered to pups, ensuring MatAb titer was not a variable impacting the results.

**Figure 4.**
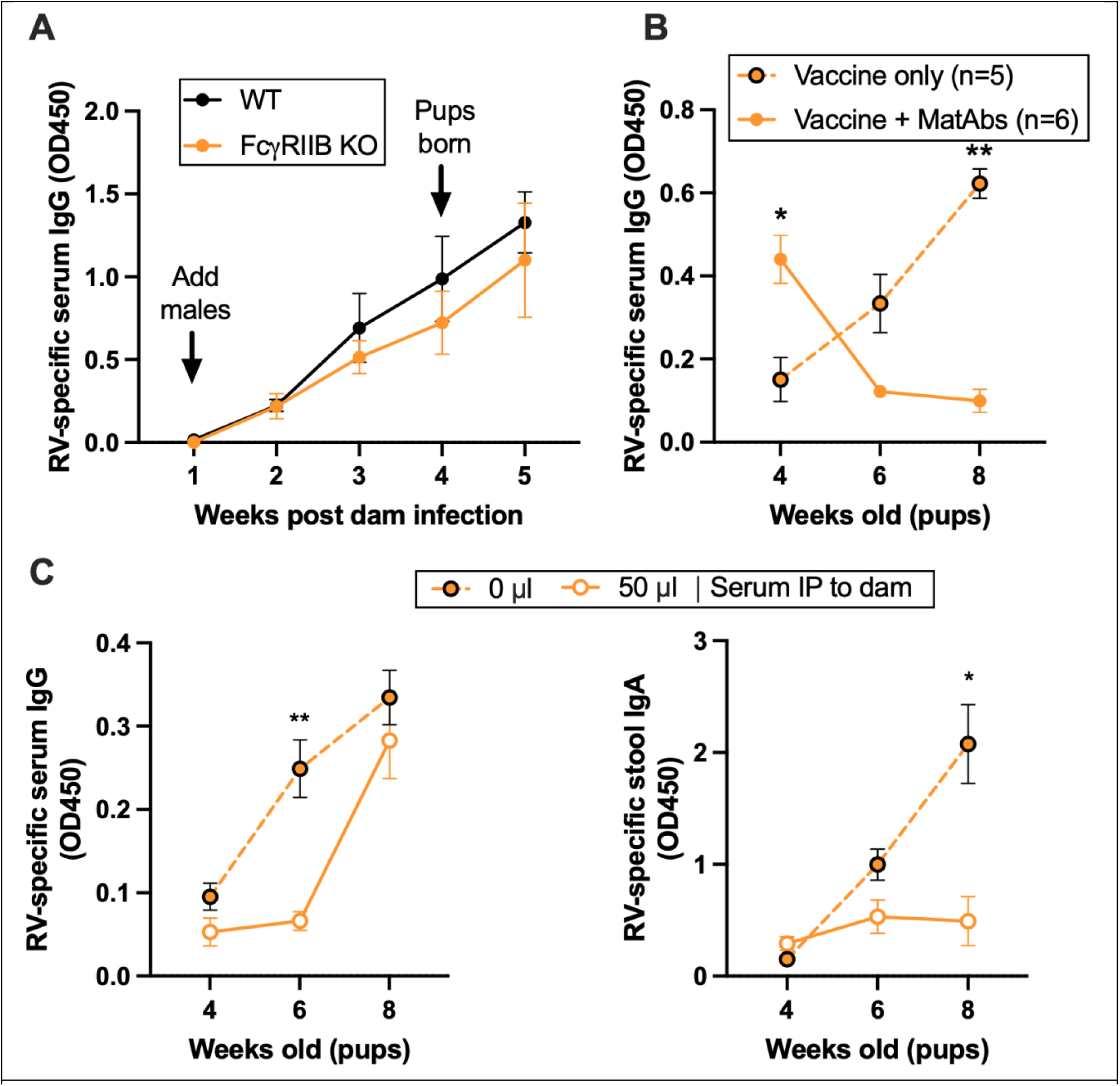
Impact of FcγRIIB signaling on maternal antibody interference. (**A**) Serum IgG responses of wild-type (WT) and FcγRIIB knockout (KO) dams following rotavirus infection as shown by ELISA. (**B**) Evaluation of seroconversion by KO pups to rotavirus vaccination in the presence / absence of maternal antibodies by serum IgG ELISA. (**C**) Serum IgG and stool IgA responses to rotavirus vaccination in pups vaccinated 24 h after dams were administered rotavirus-positive serum intraperitoneally (IP). Statistical significance was determined by two-way ANOVA with repeated measures. Significant Tukey’s adjusted pair-wise comparisons between WT and KO (**A**) and vaccination in the absence or presence of MatAbs or passively transferred serum are shown (* *p* ≤ 0.05, ** *p* < 0.01).

Figure 4B shows the seroconversion of KO pups vaccinated with rotavirus in the presence or absence of MatAbs. MatAb interference is clear in the group of pups receiving MatAbs from naturally infected dams, with pup serum IgG titers only increasing in the absence of MatAbs. This indicates that FcγRIIB signaling is not essential for MatAb interference in this experimental set up. We next asked whether FcγRIIB could be playing a more important role when MatAb titer was very low. As we had already established that the most reproducible approach to ensuring low titers of MatAbs was passive transfer of pooled serum from seropositive mice, we repeated this experimental strategy in FcγRIIB KO mice. We selected the same low dose of immune sera (50 μl) that induced interference in WT mice (Figure 3D-F), and this was administered to lactating dams 24 hours before pups were vaccinated at 7 days old. These pups were then longitudinally sampled post-weaning for analysis of their antibody responses. As shown in Figure 4C, MatAb interference was observed when measuring serum IgG responses in 6-week-old mice, but this difference was no longer apparent when mice were 8 weeks of age. However, when stool IgA responses were quantified by ELISA, there was no evidence of pup IgA production in the presence of MatAbs at any time point. This suggests that FcγRIIB may be involved in driving IgG-mediated MatAb interference at very low doses of MatAbs delivered via milk, but the mechanisms by which pup IgA responses are induced can be separated. As IgA cannot engage FcγRIIB, it is intriguing to consider if this reflects differences in how MatAb isotypes mediate interference.

### Maternal antibodies mediate vaccine clearance

A leading hypothesis for the mechanism by which MatAbs interfere with vaccination response in neonates is clearance or neutralization of the vaccine particles (1, 19, 26). Currently licensed rotavirus vaccines in human infants are all live-attenuated vaccines, and replication in the neonatal gastrointestinal tract is understood to be important for inducing an optimum vaccine response (28, 29). To investigate whether rotavirus-specific MatAbs altered replication of the murine live-attenuated vaccine, we sought to quantify vaccine replication in the pup gastrointestinal tract. Two litters of 7-day old mice were vaccinated in the presence of MatAbs, and two litters from naïve dams were vaccinated. Stool samples were collected once daily from every pup for 7 days after vaccination. Samples from each litter were pooled to generate sufficient material for analysis. qPCR was performed on nucleic acid extracted from stool samples and showed that replication of the viral vaccine was substantially reduced in the presence of MatAbs (Figure 5A). This suggests MatAbs in the intestinal mucosa are binding to viral vaccine particles and blocking replication.

**Figure 5.**
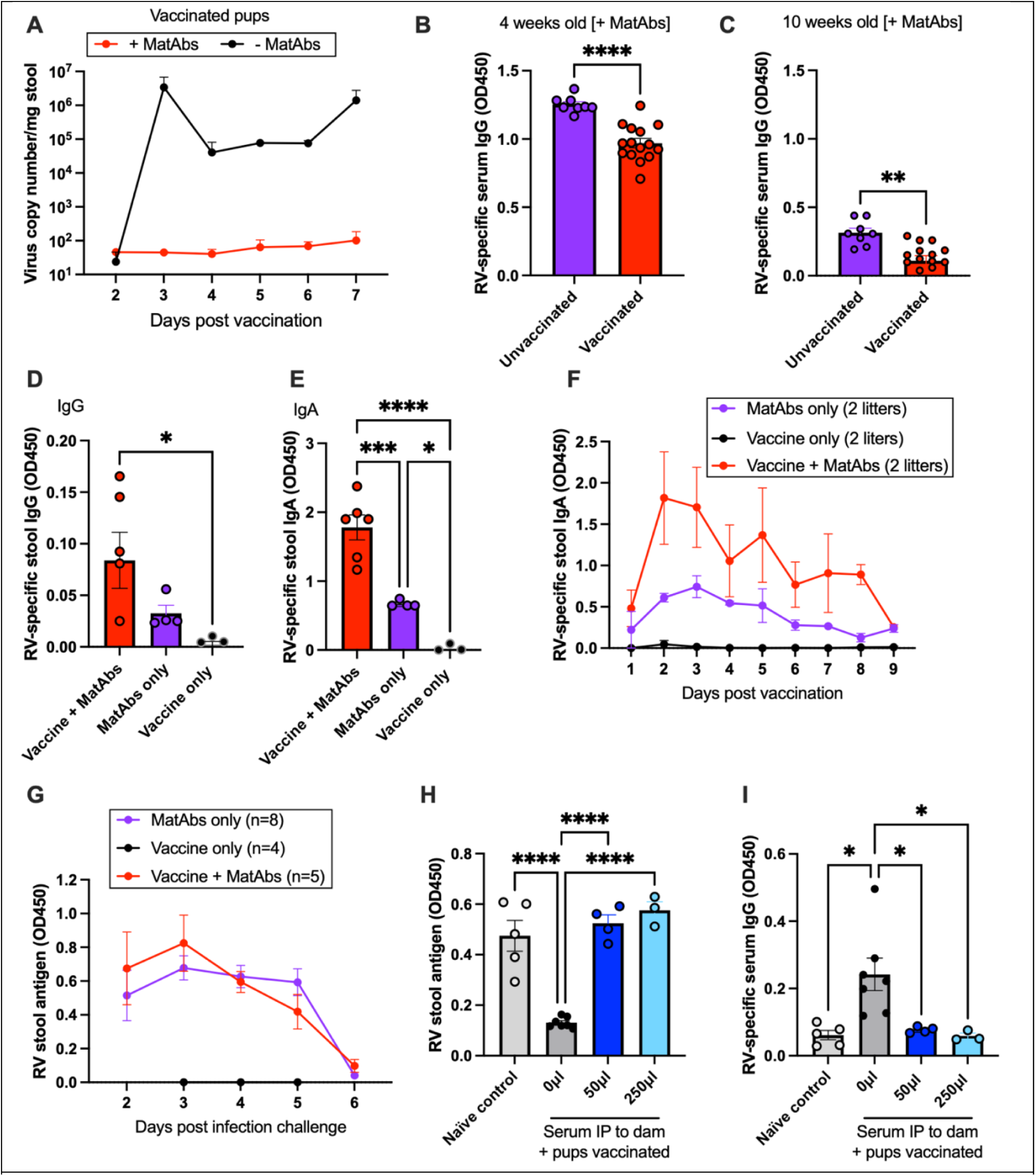
Combined clearance of vaccine and maternal antibodies, with no long-term effects on immunity. (**A**) qPCR of rotavirus in stool samples collected after vaccination in pups with and without MatAbs (2 litters each condition, samples from a litter pooled). Rotavirus-specific IgG was quantified by ELISA in mice receiving MatAbs that did (2 litters, n=15) or did not (1 litter, n=8) receive a vaccine at 7 days old, in serum samples collected at 4 weeks (**B**) or 10 weeks (**C**) of age. IgG (**D**) or IgA (**E**) were quantified in stool samples by ELISA in 9 day old pups that did/did not receive rotavirus vaccination. (**F**) Stool samples were pooled from each litter (2 litters per condition) and IgA quantified by ELISA. (**G**-**I**) Adult mice born +/- MatAbs were challenged with rotavirus when 10-12 weeks of age; rotavirus shedding was quantified by viral antigen ELISA in (**G**) mice born to naturally infected dams, and (**H**) mice born to dams receiving MatAbs IP. (**I**) Rotavirus-specific IgG induced or boosted by challenge infection as quantified 2 weeks post infection. For all graphs, error bars represent standard error. Statistical significance was determined by unpaired two-tailed t-test (**B** and **C**), one-way ANOVA (**D**, **E**, **F**, **H** and **I**), or two-way ANOVA with repeated measures (**F** and **G**). Significant Tukey’s adjusted pair-wise comparisons (**F** and **G**) and Bonferroni-corrected pair-wise comparisons (**D**, **E**, **H** and **I**) are shown (* *p* ≤ 0.05; ** *p* < 0.01, *** *p* < 0.001, **** *p* < 0.0001).

In addition to reduced vaccine replication in the presence of MatAbs, we also observed a surprising reduction in total circulating IgG MatAbs in vaccinated pups compared to unvaccinated controls (Figure 5B/C). This difference is highly significant in 4-week-old pups (*p* < 0.0001), and still maintained when pups are 10 weeks of age (*p* = 0.0014). We propose this data supports the hypothesis that vaccine clearance is a key mechanism by which interference occurs; when MatAbs bind to vaccine particles in the intestine this results in clearance of MatAb-vaccine complex. This process can be described as ‘antigen consumption’ (30), and in turn this alters the concentration gradient between MatAbs in the circulation and MatAbs in the intestinal mucosa such that more MatAbs are drawn out of circulation.

Next, we questioned what the fates of rotavirus-specific IgG MatAbs were once they had bound to their target. Immune complexes in circulation are typically cleared by innate immune cells such as phagocytes, but in the gastrointestinal tract we asked if MatAb-vaccine complexes would simply be shed in stool. To investigate this, we quantified MatAbs present in stool collected from the intestinal tract of pups at 9 days old, with or without vaccination at 7 days of age. As shown in Figure 5D, there was a trend towards more IgG MatAbs detected in stool samples if the pups were vaccinated, but this was not statistically significant. We also quantified IgA in the same stool samples, which would represent MatAbs of breast milk origin (Figure 5E). Interestingly, a significant difference between IgA MatAbs detected in stool in vaccinated and unvaccinated pups was evident. To evaluate whether this vaccination-associated increase in MatAb shedding continued in the week after vaccination, IgA was quantified in stool samples pooled from 4 litters of vaccinated pups (+/- MatAbs, same pups as in Figure 5A) and an additional 2 litters of unvaccinated pups. As shown in Figure 5F, there was a trend towards higher shedding of IgA MatAbs in the stool of pups that were vaccinated compared to those that were not, especially in the first few days after vaccination (IgG levels were below the lower limit of detection). We suggest that this provides evidence that vaccination enhances clearance of both IgG and IgA MatAbs in part via the gastrointestinal tract, although we cannot rule out increased delivery of MatAbs due to IFNγ-induced upregulation of pIgR in the mammary gland.

We reasoned that if vaccine clearance was the primary mechanism of interference, then the response to rotavirus infection when MatAbs had waned would be the same in naïve versus vaccinated + MatAb mice. To investigate this, we challenged naïve and vaccinated mice with a high dose of rotavirus (10 FFU) when MatAbs had waned at 10-12 weeks of age. Stool samples were collected daily from each mouse for quantification of virus shedding by viral-antigen ELISA, then terminal serum samples were collected 2 weeks later. As shown in Figure 5G, virus shedding in mice that had received MatAbs as pups from naturally infected dams was identical between pups vaccinated at 7 days of age and unvaccinated mice. To confirm our findings, we also challenged mice that were born to dams who received MatAbs by passive transfer (as in Figure 3E/F). We found that virus shedding of stool antigen was the same in naïve control adult mice as compared to two litters of mice that were vaccinated in the presence of MatAbs, as shown 4 days post infection in Figure 5H. Examination of antibody responses 2 weeks after virus challenge identified comparable serum IgG titers induced in naïve and MatAb groups (Figure 5I), and no differences in stool IgA across all groups (Figure S2B).

### Single cell analysis reveals interference with global anti-viral response

Whilst our results had so far proven the ability of MatAbs to interfere with germinal center formation and antibody production, the effect of MatAbs on individual immune cell responses were not well defined. To address this, we comprehensively profiled immune diversity at the single cell level of the draining mesenteric lymph node (MLN). This aimed to identify genes and cell types that were altered by the presence of MatAbs during rotavirus vaccination (Figure 6A). We used the 10X platform to perform scRNA-seq to profile cells and cellranger count outputs were imported into the Seurat pipeline to generate a landscape for cell-type annotation and exploratory analysis. After excluding low-quality cells, potential doublets, and non-lymphoid cells, we profiled a total of 31,791 cells from the vaccine + MatAbs (15,334 across 2 biological replicates) and vaccine only (16,457 across 2 biological replicates) conditions. Cluster analysis of all cells identified 18 distinct clusters and we used Seurat to examine canonical markers and identify marker genes to annotate the majority of the clusters (Figure 6B/C). All cell types were represented in all 4 samples, and we compared the proportions of each cell type between vaccine + MatAbs and vaccine only groups (Figure 6D). B cells were the most abundant cell type (39 vs 36% for vaccine + MatAb and vaccine only, respectively, *p* = 0.5) in all samples. Differences in the immune cells altered by condition were limited, although differences were identified for activated CD8 T cells (1.2 vs 3.2% for vaccine + MatAb and vaccine only, *p* = 0.06), cytotoxic CD8 T cells (1.8 vs 2.4% for vaccine + MatAb and vaccine only, *p* = 0.05) and pDC (0.17 vs. 0.36 % for vaccine + MatAb and vaccine only, *p* = 0.03). Marginally decreased activated and cytotoxic CD8 T cells at 10 days post-vaccination is consistent with the absence of viral replication and shedding in MatAb mice following vaccination.

**Figure 6.**
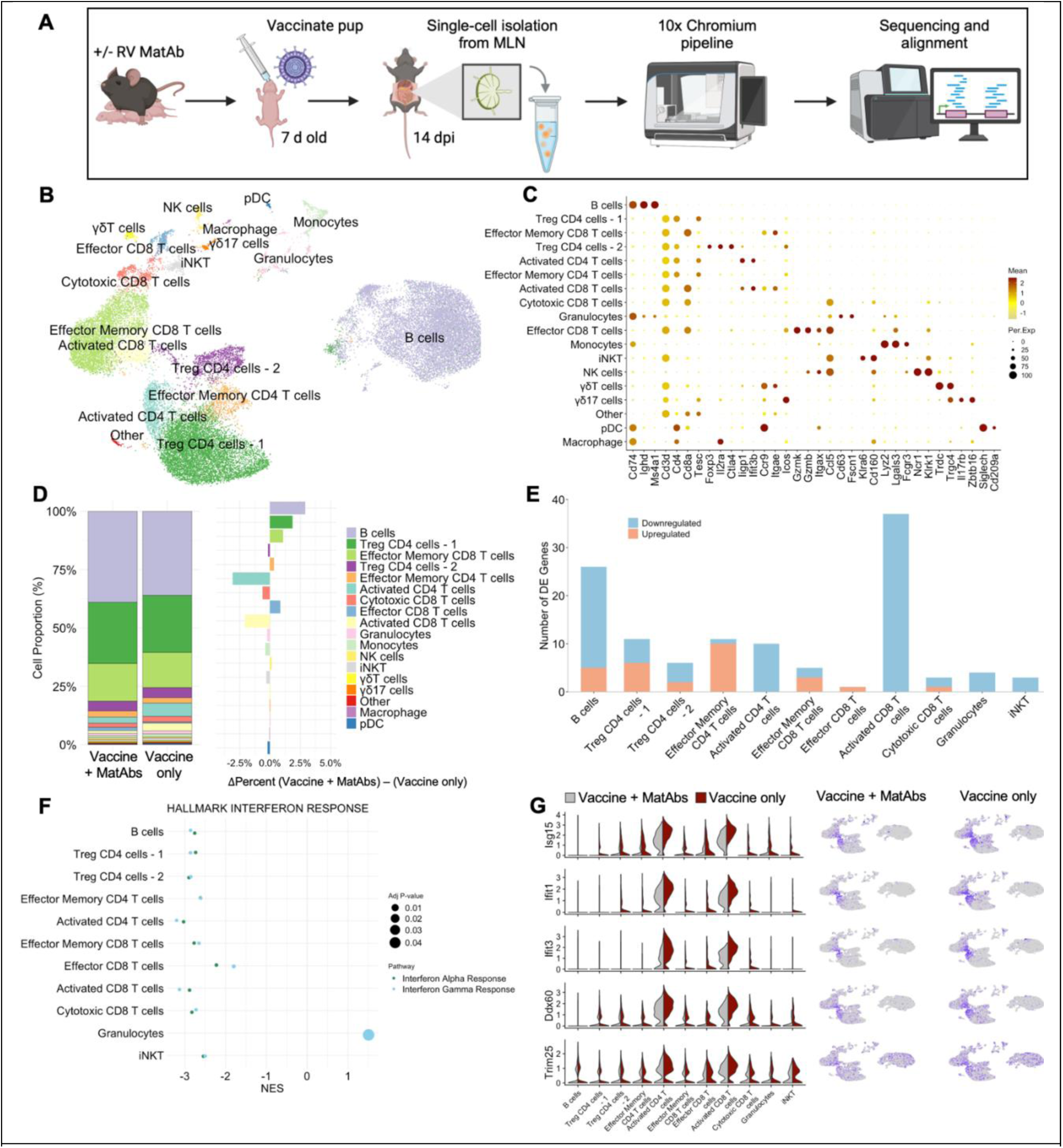
Single-cell transcriptomics during maternal antibody interference. (**A**) Schematic diagram of experimental protocol. (**B**) Integrated uniform manifold approximation and projection (UMAP) for all samples. Individual cells plotted. (**C**) Relative expression of canonical and marker genes (x axis) for immune cells across clusters (y axis); dots indicate average expression and percentage of cells detected with expression (color and size, respectively). (**D**) average proportion of cell types in vaccine + MatAb and vaccine only samples (left) and their change in proportion between vaccine + MatAb and vaccine only (right). (**E**) Counts of differentially expressed (DE) genes (y axis) per cell type, comparing vaccine + MatAb and vaccine only with absolute log2 fold change >1 and adjusted p-value ≤0.05. (**F**) GSEA results showing downregulation of hallmark interferon alpha and gamma response in major clusters. Dots are sized to denote significance; x axis indicates NES. (**G**) Distribution and expression of selected interferon stimulated genes.

To further characterize the immune cell transcriptome of the MLN we used a psuedobulk approach to aggregate counts across cells per cell type within sample. This approach to treat samples as independent observations has been shown to account for within-sample correlation and outperform Seurat (31). Given the robust failure of pups to seroconvert following vaccination in the presence of MatAbs we were specifically interested in identifying transcriptome changes in the MatAb group and used a contrast to compare vaccine + MatAb and vaccine only in differential expression analysis. We hypothesized naïve pups would mount the requisite immune response to vaccination that would result in differentiation of antibody secreting cells and any changes observed in the MatAb group would be associated with the failure to seroconvert. We further hypothesized that if vaccine clearance is a predominant mechanism of interference, there would be limited evidence of immune activation in lymphocytes in the draining lymph node. Our approach identified individual genes with expression differences between vaccine + MatAb and vaccine only for cell types with greater than 282 cells per cluster but not for less abundant cell types (Figure 6E). Of note, more downregulated genes were identified in the MatAb group across altered cell types.

To characterize the coordinated shifts in gene expression reflecting changes in pathway activation between vaccine + MatAb and vaccine only, we performed gene set enrichment analysis (GSEA) on each cluster in which we detected differential gene expression (Figure S3A). The most substantial finding from this analysis was the downregulation of interferon responses, in particular, gene sets associated with the hallmark interferon alpha and interferon gamma response were downregulated in nearly all cell types of interest (Figure 6F). We further investigated a subset of interferon stimulated genes (ISGs) previously associated with rotavirus infection in enterocytes in mice (32). As expected, the expression of selected ISGs was lower in vaccine + MatAb compared to vaccine only (Figure 6G). These results demonstrate a robust and global decrease in the transcriptome that drives response to vaccination.

To further investigate the effect of MatAbs on B cell populations in the MLN, we focused on B cell lineage in a separate analysis. Re-clustering of B cells revealed some cells with both B cell and T cell lineage markers (*Cd3d*), which were excluded, and a small cluster (116 cells) driven by an uncharacterized transcript (*Gm53771*) that was removed from the top 10 VariableFeatures before re-clustering. In the analysis of B cells, two clusters displayed a naïve phenotype and were annotated as naïve - 1 and naïve - 2 (Figure 7A/B). Activated B cells were characterized by increased expression of *Stat1*, *Ifit3* and other ISGs (data not shown). The detectable expression of the joining chain of multimeric antibodies (*Jchain*), the marker for somatic hypermutation and class switch recombination (*Aicda*), and proliferation markers (*Stmn1, Mki67*) clearly identified a cluster of plasma and germinal center (GC) B cells undergoing active proliferation and differentiation into antibody secreting cells (Figure 7B). The proportion of B cell types was not altered except for a marginal increase in the naïve - 1 cluster (*p* = 0.03) in vaccine + MatAb compared to vaccine only, 68.1 vs. 63.8%, respectively, and a decrease (*p* = 0.03) in plasma and GC B cells in vaccine + MatAb compared to vaccine only, 0.64 vs. 1.65%, respectively (Figure 7C). Along with the decrease in plasma and GC B cells, it followed that counts for differentiation markers for this population were lower or undetected but that *Ighd* was greater in vaccine + MatAb compared to vaccine only (Figure 7D). Taken together, these data confirm that Ig class switching, cell proliferation, and initiation of plasma cell differentiation were limited in the presence of MatAbs. Differential expression and GSEA further revealed an altered B cell transcriptome (Figure 7E/F, Figure S3E), including a downregulated interferon response (Figure 7G).

**Figure 7.**
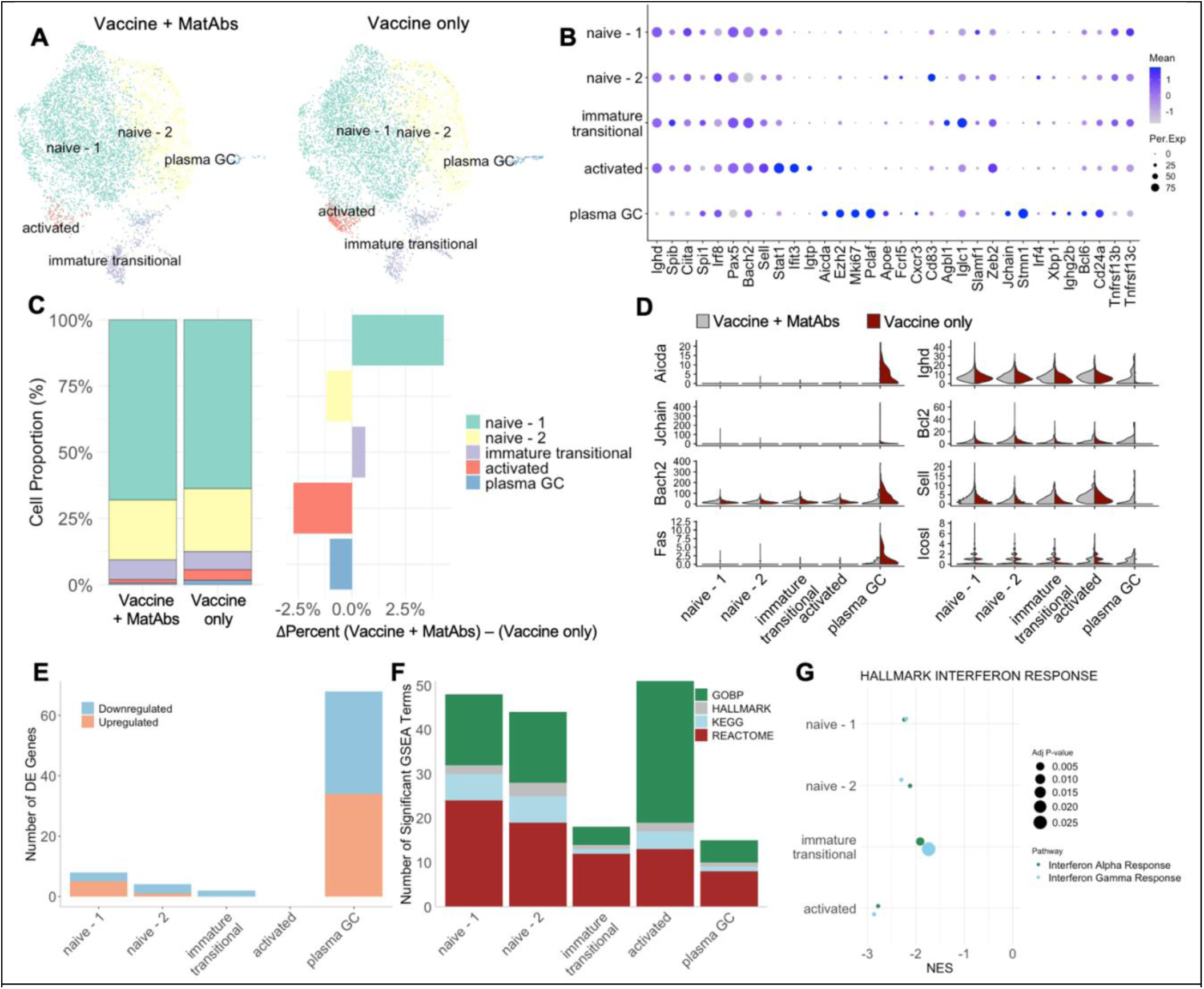
B cell transcriptomics during maternal antibody interference. (**A**) Integrated uniform manifold approximation and projection (UMAP) for vaccine + MatAb and vaccine only samples. Individual cells plotted. (**B**) Relative expression of canonical and marker genes (x axis) for immune cells across clusters (y axis); dots indicate average expression and percentage of cells detected with expression (color and size, respectively). (**C**) average proportion of cell types in vaccine + MatAb and vaccine only samples (left) and their change in proportion between vaccine + MatAb and vaccine only (right). (**D**) Distribution and counts of selected B cell differentiation markers. (**E**) Counts of differentially expressed (DE) genes (y axis) per B cell cluster, comparing vaccine + MatAb to vaccine only with absolute log2 fold change >1 and adjusted p-value ≤0.05. (**F**) Counts of the strongest significantly enriched gene sets with an absolute normalized enrichment score (NES) >2 across B cell clusters. (**G**) GSEA results showing downregulation of hallmark interferon alpha response and hallmark interferon gamma response in B cell clusters. Dots are sized to denote significance; x axis indicates NES.

In conclusion, we observed a blunted interferon response and diminished B cell activation and differentiation in the presence of MatAbs. These results support the hypothesis of reduced antigen encounter and vaccine clearance by MatAbs that blunted the antibody response to vaccination.

### Methods to overcome maternal antibody interference

The predominant strategy used to reduce the negative effect of MatAbs on human infant responses to vaccines is to delay vaccination until MatAbs are expected to have sufficiently waned (1, 33). We asked whether delaying vaccination would permit pups to seroconvert to rotavirus vaccination in our experimental model. Instead of vaccinating at 7 days old (modelling 8 weeks old in human infants), we vaccinated at weaning (3 weeks old) or at 4 weeks old when breast milk MatAbs are not detectable in the gastrointestinal tract (data not shown). As shown in Figure 8A, delaying vaccination in our system still resulted in robust MatAb interference. This indicates that circulating antibodies in the pup alone are sufficient to interfere with immunity in mice.

**Figure 8.**
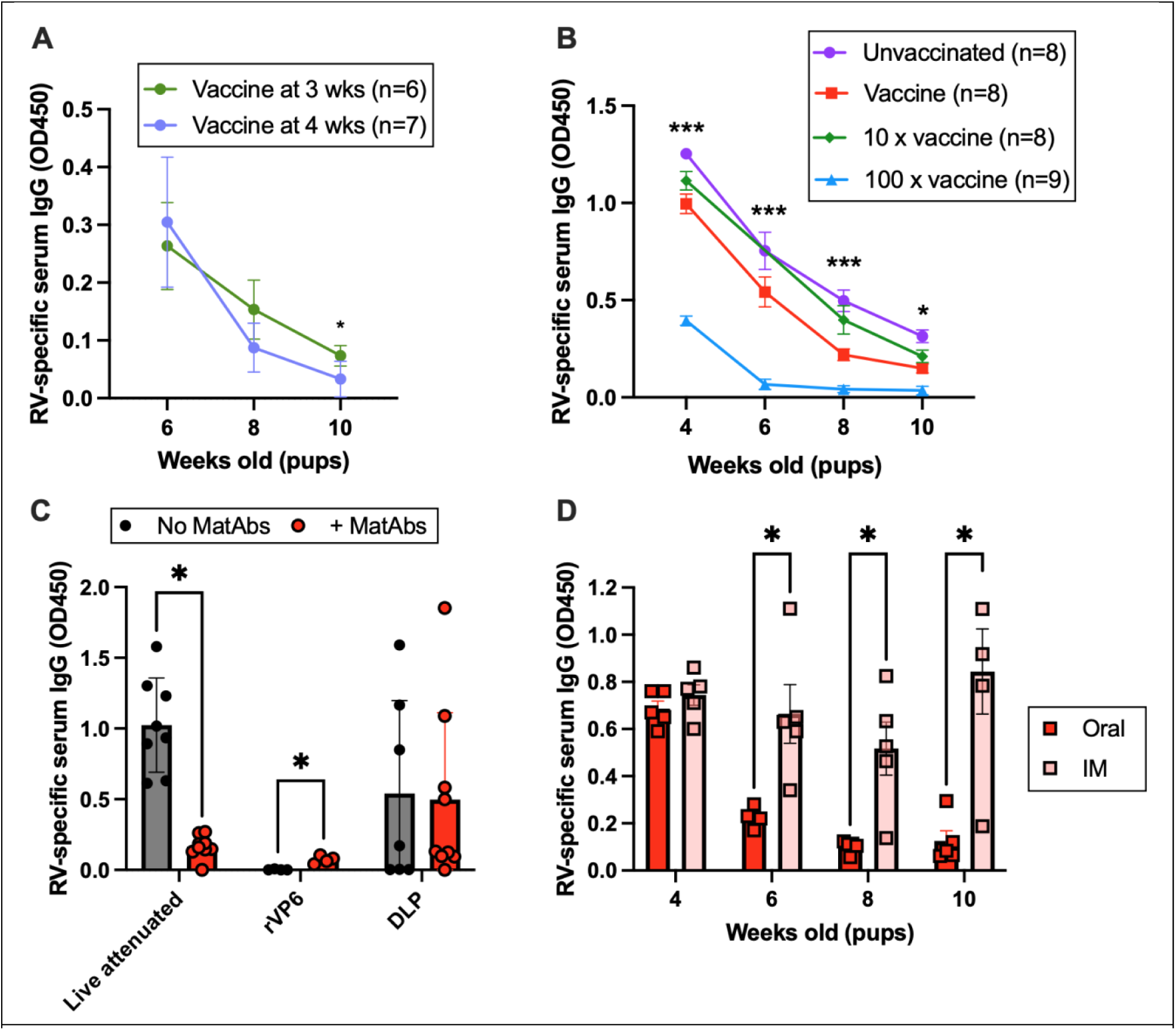
Overcoming maternal antibody interference. (**A**) Rotavirus-specific IgG in the serum of pups with MatAbs that were vaccinated at 3 or 4 weeks old. (**B**) Serum IgG ELISA of litters of pups that received MatAbs, which were unvaccinated or vaccinated with increasing doses of the rotavirus vaccine at 7 days old. (**C**) Pups born to seronegative (no MatAbs) or seropositive (+ MatAbs) dams were vaccinated at 7 days old with the standard live attenuated vaccine, recombinant VP6, or purified DLPs. Presented is serum rotavirus-specific IgG when pups were 10 weeks old. (**D**) Longitudinal serum IgG analysis of pups vaccinated in the presence of MatAbs with DLPs delivered orally with cholera toxin adjuvant, or administered intramuscularly (IM) with Addavax adjuvant. Statistical significance was determined by two-way ANOVA with repeated measures (**A**, **B** and **C**) and unpaired two-tailed t-tests (**C**). Significant Bonferroni-corrected pair-wise comparisons (**A**, **C** and **D**) and Tukey’s adjusted pair-wise comparisons between vaccine and 100x vaccine (**B**) are shown (* *p* ≤ 0.05; ** *p* < 0.01, *** *p* < 0.001).

Given a clear role for MatAb-mediated vaccine clearance in our model, we predicted that there must be a vaccine dose whereby MatAbs are out-competed and seroconversion can take place. Keeping MatAb titers the same, we immunized different litters of pups with increasing doses of vaccine, and monitored for seroconversion after weaning. Figure 8B shows that increasing vaccine dose simply increased the rate at which MatAb waning occurred. We were unable to identify a threshold over which vaccine dose induced seroconversion.

Next, we sought to determine if a different vaccine type could overcome MatAb interference. To study this, we chose to compare non-replicating subunit vaccines with the standard live-attenuated vaccine via the same oral delivery route. We immunized litters of pups with a single rotavirus protein, the inner capsid protein VP6, in the presence or absence of MatAbs. VP6 is known to be highly conserved and highly immunogenic, and has been considered as a vaccine target by multiple groups (34). We studied two different formats of VP6; either purified recombinant protein (kind gift from James Crowe), or in the form of a non-infectious double-layered particle (DLP), purified from infected cells by ultracentrifugation as previously described (35). Pups were orally vaccinated at 7 days old with 10 µg of each preparation, and blood samples collected at 4-10 weeks of age as standard. As shown in Figure 8C, when week 10 rotavirus-specific IgG was quantified by ELISA, recombinant VP6 was not immunogenic in any pups. In contrast, DLPs induced seroconversion in pups to a comparable level in pups with or without MatAbs. This indicated that MatAbs were not able to sufficiently clear DLPs and evade activation of neonatal B cells. However, DLPs alone were not as immunogenic in all pups as the live-attenuated vaccine approach.

Finally, we asked whether the route of vaccine delivery could be modified to improve vaccine efficacy in the presence of MatAbs. Vaccine administration via different routes has previously been shown to be useful in reducing MatAb interference (36, 37). We chose to compare intramuscular (IM) delivery with oral delivery, predicting that IM administration could improve vaccine efficacy. For this experiment we also included adjuvants as IM administration alone is known to have poor immunogenicity (38). DLPs were mixed with Addavax (InvivoGen, MF59-like) for IM delivery, or cholera toxin B (CTB, Sigma Aldrich) for oral delivery as Addavax is not recommended for oral administration. Figure 8D presents longitudinal rotavirus-specific IgG responses in mice vaccinated in the presence of MatAbs. Whilst oral vaccine delivery with CTB was less immunogenic than expected, 80% (4/5) of pups in a litter overcame MatAb interference by IM administration of DLPs to mount a robust IgG response.

## Discussion

Inducing protective immune responses in neonates is a major issue due to the dual challenge of immaturity of the neonatal immune system and MatAb inference (39, 40). However, despite the phenomenon of MatAb interference first being reported in 1958 (5), the fundamental mechanisms responsible for this problem have been difficult to unravel. The need for suitable experimental models to study the effects of MatAbs on neonatal immune responses has long been sought after (1, 40). Here we have presented our solution to understanding how an orally-delivered live-attenuated vaccine can be blocked by MatAbs. We have focused on rotavirus as a significant global health concern, and demonstrated that MatAbs are highly efficient at clearing rotavirus vaccines from the gastrointestinal tract in our mouse model. We propose that this contributes to the poor seroconversion observed in low-and middle-income countries to current rotavirus vaccines.

A small number of earlier studies have examined the effect of MatAbs on different types of rotavirus vaccines in mice (36, 37, 41). Johansson et al, 2008, used triple-layered virus-like particles as the immunogen and observed MatAb interference with seroconversion in offspring. Yang et al, 2019, used live murine rotavirus of the EDIM strain as the vaccine and also demonstrated MatAb interference. The robust MatAb interference observed in these studies and our model is unexpectedly distinct from results presented in a recent pre-print by Muleta et al (42). Interestingly, in that study it was reported that humoral immunity was still induced in the presence of protective MatAbs, albeit delayed until after weaning. This is in contrast to our findings where no humoral immunity was observed for at least seven weeks after weaning. A key difference between these studies is that Muleta et al challenged pups with a high dose of the virulent murine rotavirus strain ECw, whereas in our study we have modelled vaccination using low doses of the attenuated murine rotavirus strain EMcN. We therefore propose that our model is more reflective of MatAb interference with rotavirus vaccination, whereas Muleta et al demonstrate interesting findings that align with potent natural infection.

A role for FcγRIIB in driving IgG-mediated interference has long been considered in a variety of different immunological contexts. Our results in an FcγRIIB KO mouse line have shown that FcγRIIB is not central to MatAb interference to rotavirus vaccination. This is in agreement with a previous study that examined immune responses to sheep erythrocytes (43). When anti-sheep erythrocyte IgG was passively transferred to mice at the same time as sheep erythrocytes, no anti-sheep erythrocyte IgG was induced in either wild-type or knockout animals. Our FcγRIIB results are contradictory to some degree with those reported using a cotton rat model to study MatAb-mediated interference with measles vaccination (21). In this study, F(ab’)2 fragments passively transferred at the same time as vaccination did not suppress neutralizing antibody production. It was concluded that this indicated that the Fc region of IgG was required for suppression. Furthermore, only IgG isotypes capable of binding to FcγRIIB mediated suppression. However, there are key caveats to this study that could explain our different conclusions. Firstly, being a rat model of vaccination, no FcγRIIB KO animals were available and results are therefore largely indirect. Secondly, whereas our model uses an orally-delivered vaccine which can replicate in the gastrointestinal tract, their model used a subcutaneously-delivered vaccine expected to replicate systemically.

Our results provide strong evidence for MatAb-mediated clearance of vaccine particles, which prevents the neonatal B cell from encountering their target. This is in agreement with another recently reported study focusing specifically on breast milk antibodies to SARS-CoV-2 in a mouse model (44). However, the fate of MatAb-vaccine complexes is uncertain. We proposed that some of the MatAb-vaccine complex would be cleared from the gastrointestinal tract in stool, and indeed we did observe increased shedding of IgA MatAb in stool samples following vaccination. However, an alternative hypothesis is that vaccination increases pIgR expression in the mammary gland, which is known to be upregulated by interferon-g (45). This would result in more IgA being delivered to the pup gastrointestinal tract. It is also possible that MatAb-vaccine complexes are cleared by antibody-mediated phagocytosis (19). We were unable to detect any differences in phagocyte populations in our single cell RNA-sequencing analysis, but this could be due to the low number of phagocytes captured, or the location in which phagocytes mediate such activity.

Our single cell RNA-sequencing analysis has provided a detailed insight into the cell population and gene expression changes that occur in MLN after live-attenuated rotavirus vaccination in the presence and absence of MatAbs. Our overarching finding was that vaccination of naïve neonatal mice induces a robust set of changes in anti-viral immune pathways, which are still readily detected two weeks post-vaccination. In the presence of MatAbs there was a significant decrease in activation of any of these pathways, which is in agreement with our more focused experimental approaches. Our study is the first to report single-cell analysis of the lymph node response to rotavirus infection, and as such provides valuable insight into the anti-viral pathways stimulated. Previous work has shown significant interferon-related pathways are induced in intestinal epithelial cells where CD45+ cells were removed from sample processing (32), so our study provides useful complimentary results.

A limitation of using a mouse model to study MatAb transfer is the biological differences in transfer mechanisms relative to humans. Whilst both humans and mice have a hemochorial placenta, making mice a good model in comparison to larger animal species with different placentation types, significantly fewer MatAbs are delivered transplacentally in mice. Whereas the human placenta permits transfer of the majority of IgG, in mice only 30% of maternal IgG is found in pups at birth (46). To make up for this reduced transfer pre-birth, mouse milk contains significantly higher titers of IgG than human milk. However, by weaning at 3 weeks of age, comparable total MatAb transfer has occurred in mice relative to humans. Throughout most of our study we therefore considered the impact of total MatAb transfer as a whole, and did not divide our conclusions into breast milk versus transplacental antibodies, nor IgG versus IgA. We had two interesting exceptions to this approach though; delivery of artificial MatAbs in the form of pooled serum administered IP to dams, and vaccination after breast feeding had ceased. These experiments allowed us to show that in mice, breast milk antibodies alone can mediate interference, and also that MatAbs in the pup circulation (which will have originated from both transplacental and milk delivery) can be sufficient to mediate interference after weaning.

A key motivator for understanding the biological mechanisms underpinning MatAb interference is the potential for rational development of vaccine strategies to overcome this issue. Although delaying vaccination for longer to permit more MatAb waning has shown potential in human infants (47), this strategy had no benefit in our mouse model, and in heterogenous human populations this certainly risks leaving some infants vulnerable for longer if MatAb delivery was impaired in any way. Increasing vaccine dose has been considered as an option for evading MatAb interference, and has shown promise in some human rotavirus clinical trials (16). However, this approach was ineffective in our model, for reasons that remain unclear but could be due to the vaccine dose:MatAb ratio. It is important to consider that increasing vaccine dose is also expected to increase the risk of side effects, which have blighted the rotavirus vaccine field since withdrawal of the first licenced rotavirus vaccine in 1999 (48). Development of subunit rotavirus vaccines for parenteral administration has been a significant avenue of research in recent years (49). Our results are in agreement with two other murine studies that demonstrated parenteral administration of rotavirus vaccines could overcome interference much more effectively than oral vaccine administration (36, 37). We demonstrated that vaccination via the intramuscular route resulted in seroconversion in the majority of mice within a cohort. These results therefore support development of a parenteral rotavirus vaccine as a strategy to minimize the negative effects of MatAbs, although the difficulties inducing mucosal immune responses via this route still require attention (50). It is clear from the results of the recent phase III vaccine trial with a parenteral non-replicating P2-VP8 vaccine that additional challenges remain (51). This parenterally delivered vaccine was proven to be less effective at preventing severe gastroenteritis than the standard oral live-attenuated vaccine. Following this, there is considerable interest in developing alternative vaccine platforms that will be effective irrespective of MatAbs. Several mRNA-based rotavirus vaccines have been reported in pre-clinical studies (52–54), which could be promising, although mRNA-based vaccines are partially affected by MatAbs for both influenza and SARS-CoV in pre-clinical vaccine studies (55, 56).

In summary, we have shown that MatAb-mediated clearance of vaccine particles is a predominant mechanism by which MatAbs interfere with the ability of the neonate to respond to oral live-attenuated rotavirus vaccines in mice. We propose that understanding this biological phenomenon will be valuable for informing future vaccine development.

## Methods

Reagent specifics are detailed in Table 1.

| Reagent | Source | Identifier |
| --- | --- | --- |
| Antibodies and Dyes |  |  |
| Anti-Ki-67-FITC | BD Pharmingen | CAT#558616; Clone: B56 |
| Anti-Mouse IgA HRP (Goat) | Bio-Rad | CAT#STAR137P |
| Anti-Mouse IgG HRP (Goat) | Sigma Aldrich | CAT#A0168 |
| Anti-IgG1a Biotin (Mouse) | BD Pharmingen | CAT#553500; Clone: 10.9 |
| Anti-Rabbit IgG HRP (Goat) | BD Pharmingen | CAT#554021 |
| APC anti-mouse CD95 (Fas) | BioLegend | CAT#152603; Clone: SA367H8 |
| Armenian Hamster anti-PD-1 | Novus Biologicals | CAT#NBP1-43110AF700 |
| CD45 (PE Rat Anti-Mouse CD45R/B220) | BD Pharmingen | CAT#553090; Clone: RA3-6B2 |
| CD45R (PerCP/Cyanine 5.5 Anti-Mouse/Human CD45R/B220) | BioLegend | CAT#103236; Clone: RA3-6B2 |
| Fc Block (TruStain FcX Anti-Mouse CD16/32) | BioLegend | CAT#101320; Clone: 93 |
| FITC anti-mouse CD185 (CXCR5) | BioLegend | CAT#145520; Clone: L138D7 |
| Fixable Viability Dye eFlour 780 (L/D) | Invitrogen | CAT#65-0865 |
| GL7 Antigen (Pacific Blue Anti-Mouse/Human) | BioLegend | CAT#144614; Clone: GL7 |
| Hoechst 33342 | Invitrogen | CAT#H3570 |
| Rotavirus Polyclonal Antibody (Rabbit) | Antibodies-online | CAT#ABIN308233 |
| Rotavirus Polyclonal Antibody (Sheep) | Invitrogen | CAT#PA1-85845 |
| Streptavidin-HRP | Mabtech | CAT#3310-9 |
| Chemicals |  |  |
| Dulbecco's Modified Eagle's Medium (DMEM) | Corning | CAT#10-013 |
| Endotoxin-Free Dulbecco's PBS (1X) | EMD Millipore | CAT#TMS-012-A |
| Fetal Bovine Serum (FBS) | Corning | CAT#28622001 |
| Oxytocin | Bimeda | CAT#1OXY015 |
| Pierce Protease Inhibitor Mini Tablets | Thermo Fisher Scientific | CAT#A32953 |
| RIPA Lysis and Extraction Buffer | Thermo Fisher Scientific | CAT#89900 |
| Sudan Black B | Ward's Science | CAT#470109-394 |
| TPCK-Treated Trypsin | Worthington Biochemical | CAT#LS003740 |
| Triton X-100 | MB Biomedicals | CAT#807423 |
| Xylenes | Thermo Fisher Scientific | CAT#L13317.AP |
| 3,3',5,5'-Tetramethylbenzidine (TMB) Membrane Peroxidase Substrate Plus | Avantor | CAT#K830 |
| Commercial Assays |  |  |
| BCA Protein Assay Kit | Thermo Fisher Scientific | CAT#23227 |
| Luna Universal One-Step RT-qPCR Kit | New England Biolabs | CAT#E3006 |
| Monarch Total RNA Miniprep Kit | New England Biolabs | CAT#T2010S |

### Sex as a biological variable

Our study examines the immunology of maternal antibodies, and therefore only breeding female mice were infected with EMcN to induce antibodies. Transfer of maternal antibodies were studied among male and female offspring. Results are therefore reported as a single cohort combining results from both sexes.

### Mice

C57BL/6 (The Jackson Laboratory), BALB/c (The Jackson Laboratory) and FcγRIIB knockout (gift from J Ravetch, Rockefeller, USA) mice were housed and bred at the Baker Institute for Animal Health, Cornell University, or the Anne McLaren Building, University of Cambridge, under specific pathogen–free conditions.

### Virus preparation, infection and vaccination

Murine rotavirus (EMcN) (22) and the heterologous rotavirus vaccine strain (Rotarix) were grown in fetal monkey kidney (MA104) cells in the presence of trypsin, as previously described (24). Virus was trypsin-activated (10 μg/ml) at 37°C for 30 minutes prior to MA104 cell infection. Virus preparations were titrated by fluorescent focus assay on MA104 cells and expressed as a number of fluorescent focus units (FFU), although replication of EMcN in vitro was limited. Virus was frozen at -80°C until use. Prior to mouse infection or vaccination virus was diluted to the appropriate titer in sterile PBS without calcium chloride and magnesium chloride. Female mice were infected orally with 10 FFU EMcN rotavirus or uninfected. Following confirmation of anti-rotavirus antibodies in the infected group, mating trios were established. At 7 days of age, pups born to dams without (naïve) or with (MatAbs) anti-rotavirus antibodies were vaccinated with a dose of 0.01 FFU EMcN by oral gavage. Purified recombinant VP6 (rVP6), provided by Dr. James Crowe (Vanderbilt Vaccine Center), and double-layered particle (DLP) purified from infected cells by ultracentrifugation, were tested as an alternative vaccine type. Addavax (InvivoGen, MF59-like) and Cholera toxin B (CTB, Sigma Aldrich) adjuvants were administered to compare an oral or intramuscular vaccine delivery.

### Milk, serum, stool and collection

To collect milk the dam was separated for approximately two hours from her litter prior to milking, then received 2 IU/kg of oxytocin intraperitoneally. Milk was aspirated from the mammary glands by manually expressing each teat. To collect serum blood was collected through the lateral saphenous vein or through cardiac puncture, then centrifuged at 6,000 × g for 5 minutes, and the serum stored at -80°C. Stool samples from neonates were pooled daily from each litter and diluted 1:10 in PBS. Stool samples from adult mice were collected and stored without pooling. Diluted stool was centrifuged at 8,000 × g for 5 minutes to remove debris, and the resulting supernatant was stored at - 80°C.

### Passive transfer of antibody mouse model

Serum from adult mice that had previously been infected with EMcN was pooled and rotavirus-specific IgG and IgA quantified by ELISA. When pups were six days old, pooled serum diluted in endotoxin-Free Dulbecco’s PBS to a total volume of 250 µl was administered to dams IP. 24 hours later each pup in the litter was infected with EMcN by oral gavage. One week after the first passive transfer of pooled serum, a second dose of pooled serum at half the original volume was administered.

### IgG, IgA and IgG1a ELISAs

Anti-rotavirus IgA, IgG and IgG1a were determined by ELISA as previously described (24, 35). Briefly, rotavirus-specific polyclonal antibody (sheep) was plated at a concentration of 5 µg/ml in PBS. Purified cell culture lysates (virus-infected lysate or mock-infected control lysate) were diluted to 10 µg/ml in PBS. All samples were tested in duplicate at a 1:200 dilution (serum), 1:50 dilution (milk), 1:1000 dilution (pup stool), or a 1:100 dilution (adult stool) in 5% milk–PBS-T. IgG antibodies were measured using a goat anti-mouse IgG secondary antibody–HRP. IgA antibodies were measured using a goat anti-mouse IgA secondary antibody–HRP. IgG1a antibodies were measured using a mouse anti- IgG1a conjugated to biotin, followed by an additional 1-hour incubation with streptavidin-HRP diluted 1:1000 in 5% milk–PBS-T. TMB detected bound antibody and the absorbance at 450 nm (OD450) was measured using a BioTek Cytation 7 Cell Imaging Multimode Reader, Gen5 Image Prime software (version 3.13).

### Antigen ELISA

Rotavirus-specific polyclonal antibody (sheep) was plated at a concentration of 5 µg/ml in PBS. Pup stool samples were tested at a 1:5000 dilution and adult stool was tested at a 1:500 dilution, then incubated at 37°C for 2 hours. The anti-rotavirus polyclonal antibody (rabbit) was diluted 1:3000 in 5% milk–PBS-T and incubated at 37°C for 1 hour, followed by the anti-rabbit IgG-HRP (goat) under the same conditions. TMB was used to detect bound antibody, then read at 450 nm (OD450) on the BioTek Cytation 7 Cell Imaging Multimode Reader, Gen5 Image Prime software (version 3.13).

### RT-qPCR

RNA was extracted from clarified stool suspensions using the Monarch Total RNA Miniprep Kit, following the manufacturer’s instructions. Extracted RNA was denatured at 95°C for 5 minutes to separate double-stranded RNA. RT-qPCR was performed using the Luna Universal One-Step RT-qPCR Kit, according to the manufacturer’s instructions and containing the NSP5 sequences: forward primer CTGCTTCAAACGATCCACTCAC 400 nM, reverse primer TGAATCCATAGACACGCC 400 nM, TaqMan probe FAM-TCAAATGCAGTTAAGACAAATGCAGACGCT-TAMRA 200 nM. Amplification was performed on a QuantStudio 3 thermocycler (Applied Biosystems) with the following cycling conditions: 55°C for 10 minutes, 95°C for 1 minute, followed by 40 × (95°C for 10 seconds, 60°C for 30 seconds). To quantify rotavirus genome copies per ml of stool supernatant, each plate included a 10-fold serial dilution of SA11 total RNA. Samples with undetectable virus were assigned a lower limit of quantification of 100 genome copy numbers. Analysis was performed using QuantStudio Design & Analysis Software (version 1.5.1).

### Mesenteric lymph node single cell suspension

Mesenteric lymph nodes (MLNs) were harvested from infected mice. Single cells were isolated by mechanical disruption through a 70 μm cell strainer. Cells were washed once with RPMI+FBS, and resuspended in PBS+1%FBS (flow cytometry) or PBS+0.04%BSA (sequencing) before filtering through a 70 μm (flow cytometry) or 35 μm cell strainer (sequencing).

### Flow cytometry (FACS)

MLN single cells were incubated with Fc Block (1:100) in staining buffer (PBS+1%FBS) for 30 minutes at 4°C, then cells were incubated with viability dye (L/D) and fluorescently conjugated antibodies targeting GC cells (GL7 Antigen and Fas) and TfH cells (PD-1 and CXCR5) in staining buffer for 30 minutes at 4°C. Cells were resuspended in 100 μl 4%PFA-PBS at 4°C for 15 minutes to fix, then resuspended in 300 μl staining buffer. Data was acquired using a BD LSRFortessa X-20 (BD Biosciences) and BD FACSDiva Software (version 9.0); analysis was performed using FlowJo (Treestar Inc, version 10.9.0).

### Single cell counting and sequencing

MLN single cells were washed once with PBS+0.04%BSA before cell counting. Single cell suspensions were run on a Chromium X instrument and libraries were prepared following the Chromium Next GEM Single Cell 3’ RNA-Seq - Dual Index Assay version 3.1 (10x Genomics, user guide CG000315, RevE) by the Cornell BRC Genomics Facility (RRID:SCR_021727).

We targeted 10,000 cells and used 13 cycles of cDNA amplification. Sample quality was confirmed using a Qubit (DNA HS kit; Thermo Fisher) to determine concentration and a Fragment Analyzer (Agilent) to confirm fragment size integrity. Libraries were sequenced on a NovaSeqX, 10B flowcell with 2 x 150bp read length.

### Data processing

Fastq files were processed by the cloud-based cellranger count (version 8.0.0) available through 10x Genomics using default parameters and a murine reference (Mouse (GRCm39) 2024-A). Matrix files (min.cells = 3, min.features = 200) from each sample were imported into R for analysis with Seurat (version 5.2.1) (57). Initial filtering removed cells that did not meet minimum quality criteria [subset(sobj, subset = nFeature_RNA > 1000 & nFeature_RNA < 6000 & percent.mt < 7.5 & log10GenesPerUMI > 0.80)]. Additional parameters were calculated for further filtering using the PercentageFeatureSet including percent.RBC (“^Hb[ab]-“), percent.platelet (“^Gypa”), and percent.CD45 (“^Ptprc”). Additional filtering was applied (percent.RBC < 0.05 & percent.platelet < 0.004 & percent.CD45 > 0.003) to remove RBC, platelets, and cells with low Ptprc expression. To limit the effect of cell cycle score and ribosomal gene expression during downstream analysis, cell cycle scores (S.score, G2M.Score) and ribosomal gene scores (R.Score) were added to the matrix using CellCycleScoring and AddModuleScore using ribosomal genes, respectively. CellCycleScoring and AddModuleScore were calculated from the “scale.data” slot after LogNormalization and ScaleData were applied to the RNA assay. Cells with R.Score less than -2 were filtered out of the object.

Normalization of UMI counts for each sample was performed using the SCTransform (version 0.4.1) (58, 59) function [SCTransform(sobj, vars.to.regress = c("percent.mt", "S.Score", "G2M.Score", "R.Score"))]. Cell doublets within each sample were determined with scDblFinder (v1.18.0) (60) and removed from the dataset. Samples were integrated using the SeuratWrappers function IntegrateLayers(method = HarmonyIntegration) (61) (with default parameters. Clustering with Seurat [FindNeighbors(sobj, reduction = “harmony”, dims = 1:50) %>% FindClusters(sobj, resolution = 0.6)] generated a total of 22 clusters. The FindMarkers function in Seurat was used to determine marker genes between clusters. Cell clusters were annotated manually using known canonical markers determined by FindMarkers. Cell demographics were calculated as cell counts per cluster per individual and normalized to the total cell counts per sample.

### Subsetting B cells

To further investigate B cell subpopulations, B cells were subsetted into a separate Seurat object and reanalyzed in the Seurat pipeline as above; however, cell cycle score was not regressed out of the matrix. Cells expressing the T cell marker Cd3d were filtered out of the B cell object before further analysis.

### Pseudobulk analysis

To account for replicate samples within each condition, a pseudobulk approach was used to investigated differentially expression genes between conditions. Raw pseudobulk counts were extracted from the Seurat object for each cluster and pseudobulk matrices were used to generate normalized counts with DEseq2 (version 1.44.0) (62). Counts were filtered to remove counts ≤10 before downstream analyses. DEseq2 was used to detect differentially expression genes between conditions [results(dds, contrast=c(‘condition’, ‘Matab’, ‘Naïve’), alpha = 0.05)]. The log2-fold change calculation from DEseq2 was used to generate a rank list of genes for gene set enrichment analysis (GSEA, (63)) using clusterProfiler (version 4.12.16) (64) to run the GSEA (method = fgsea) (65) after filtering out genes with low coverage. For example, the rank list for each cluster was filtered to retain the top quartiles of genes based on the median normalized counts across all samples. The Hallmark, C2:CP and C5 catalogs from the MsigDB database (version 7.5.1) (63) were used for enrichment tests.

### Histological and immunofluorescence analysis

Tissues were paraffin-embedded and sectioned for mounting on slides. Staining for hematoxylin and eosin (H&E) was performed on MLNs. For immunofluorescence slides were rehydrated in xylene, ethanol, and PBS, then antigen retrieval was performed by heating with citrate buffer. Tissue sections were permeabilized with 2% Triton X-100 in PBS, then washed three times with PBS-T prior to blocking with 5% BSA in PBS for 30mins. Slides were washed three times in PBS-T, then fluorescently-conjugated antibodies (Ki-67 FITC, and B220-PE) or Hoechst, in antibody dilution buffer (2% BSA in PBS) were incubated overnight at 4°C. Slides were washed three times with PBS-T, then blocked with sudan black solution (0.05% sudan black in 100% ethanol) to reduce autofluorescence. Slides were finally washed three times with PBS-T before mounting medium was used to apply a cover slip. Once dry, slides were imaged using Cytation 7 (Biotek Aligent).

### Statistics

Statistical analysis was performed using GraphPad Prism (version 10.3.1) and R (version 4.4.1). Immune response outcomes for two groups were analyzed by unpaired two-tailed t-tests. Dependent outcomes reported for three or more groups were analyzed by one-way ANOVA and pair- wise comparisons reported with Tukey’s adjustment for multiple comparisons. Outcomes reported over time were analyzed by two-way ANOVA with repeated measures with Bonferroni’s correction for multiple comparisons for two groups and Tukey’s adjustment for multiple comparisons for three groups. Statistical differences were considered significant at p-values ≤ 0.05 for all comparisons. Error bars indicated the standard error of the mean.

### Study approval

All animal studies conducted at Cornell University were approved by the Cornell University Institutional Animal Care and Use Committee (IACUC), Protocol 2022-0152. All animal studies conducted at the University of Cambridge UBS Anne McLaren facility were performed in accordance with UK Home Office guidelines and were approved by the University of Cambridge Animal Welfare and Ethical Review Board (PPL number PP6782732).

## Data availability

(1) Underlying data summarized in figures is available as supplemental information.
(2) Single-cell RNA-seq data will be deposited at GEO and publicly available as of the date of publication.
(3) Any additional information required to reanalyse the data reported in this paper is available from the corresponding author upon request.

## Acknowledgements

We thank the Center for Animal Resources and Education (CARE) team at Cornell, and the Anne McLaren Building team at the University of Cambridge for providing support with animal studies. We thank Jeffery Ravetch for the gift of FcγRIIB KO mice, and James Crowe for the gift of rVP6. We also thank Gordon Dougan, Sallie Permar and Deb Fowell for helpful discussions, and John Parker for manuscript review.

---

**Supplementary Figure 1.**
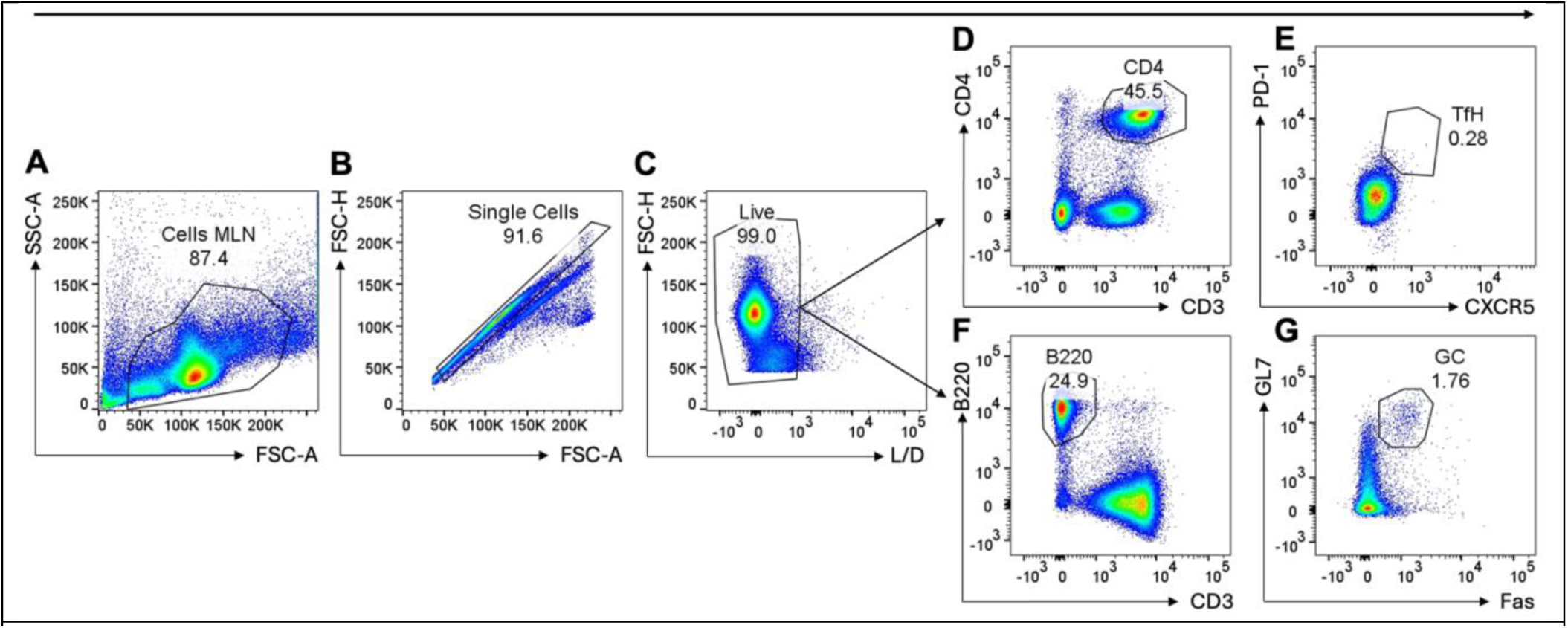
Gating strategy for identification of Tfh cells and GC B cells by flow cytometry. (**A**) Plot for forward versus side scatter, and mesenteric lymph node (MLN) cells gate. (**B**) Plot of forward scatter area as a function of forward scatter height within the MLN cells gate, and resulting single cells gate. (**C**) Viability dye (L/D) exclusion plot within the singlet gate, and resulting live cell gate. (**D**) Gating of CD4 cells, then gating of T follicular helper (Tfh) cells (**E**), identified by PD-1 and CXCR5 within the CD4 gate. (**F**) Gating of B220+ B cells, then gating of germinal center (GC) B cells (**G**), identified by Fas and GL7 cells within the B220+ gate.

**Supplementary Figure 2.**
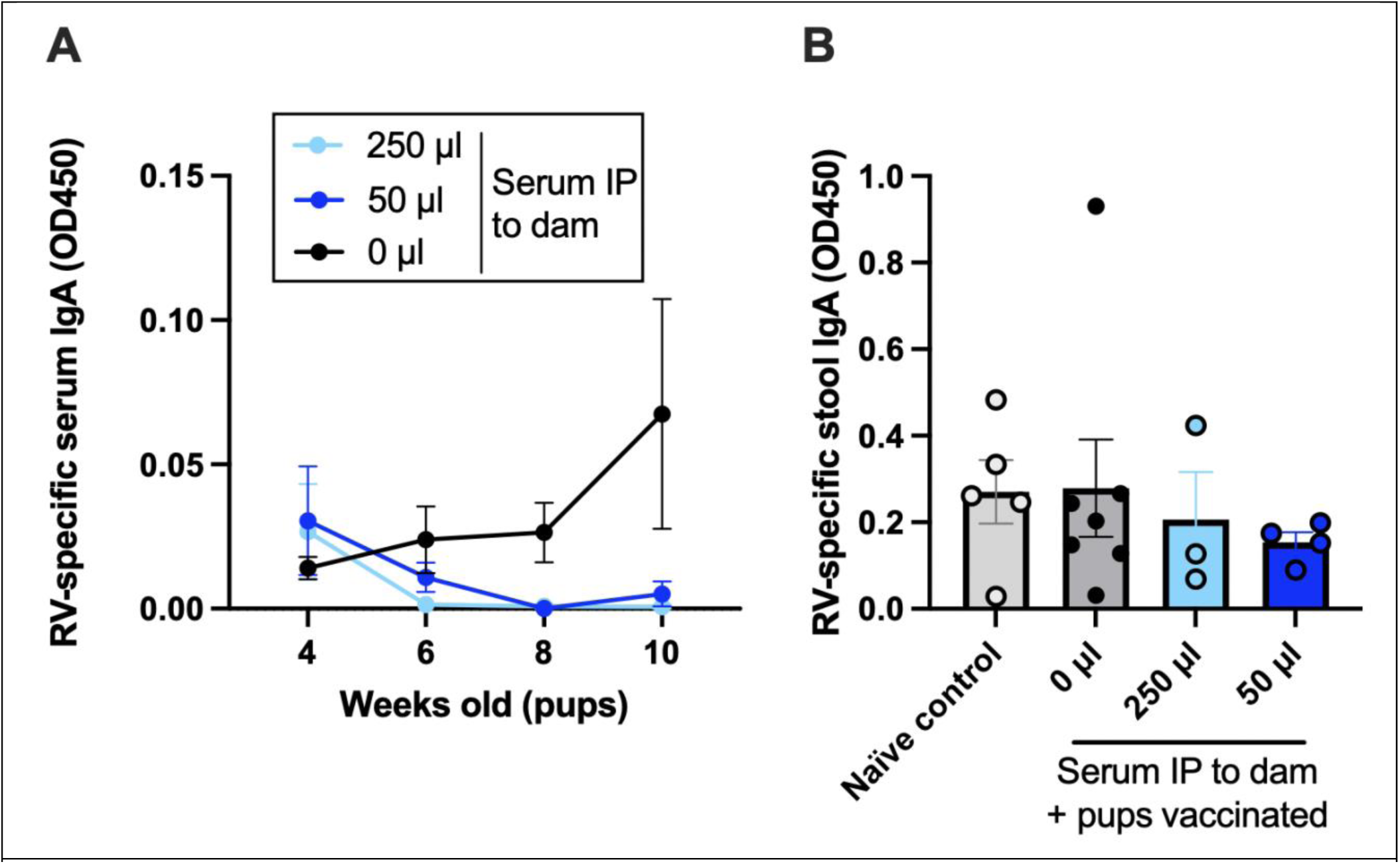
Quantification of IgA in mouse stool and serum samples in the presence or absence of MatAbs. (**A**) Longitudinal serum IgA titers following vaccination at 7 d old with different volumes of rotavirus-seropositive serum passively transferred to dams. (**B**) Stool IgA titers in adult mice from experiment (**A**) 14 days after challenge with rotavirus.

**Supplementary Figure 3.**
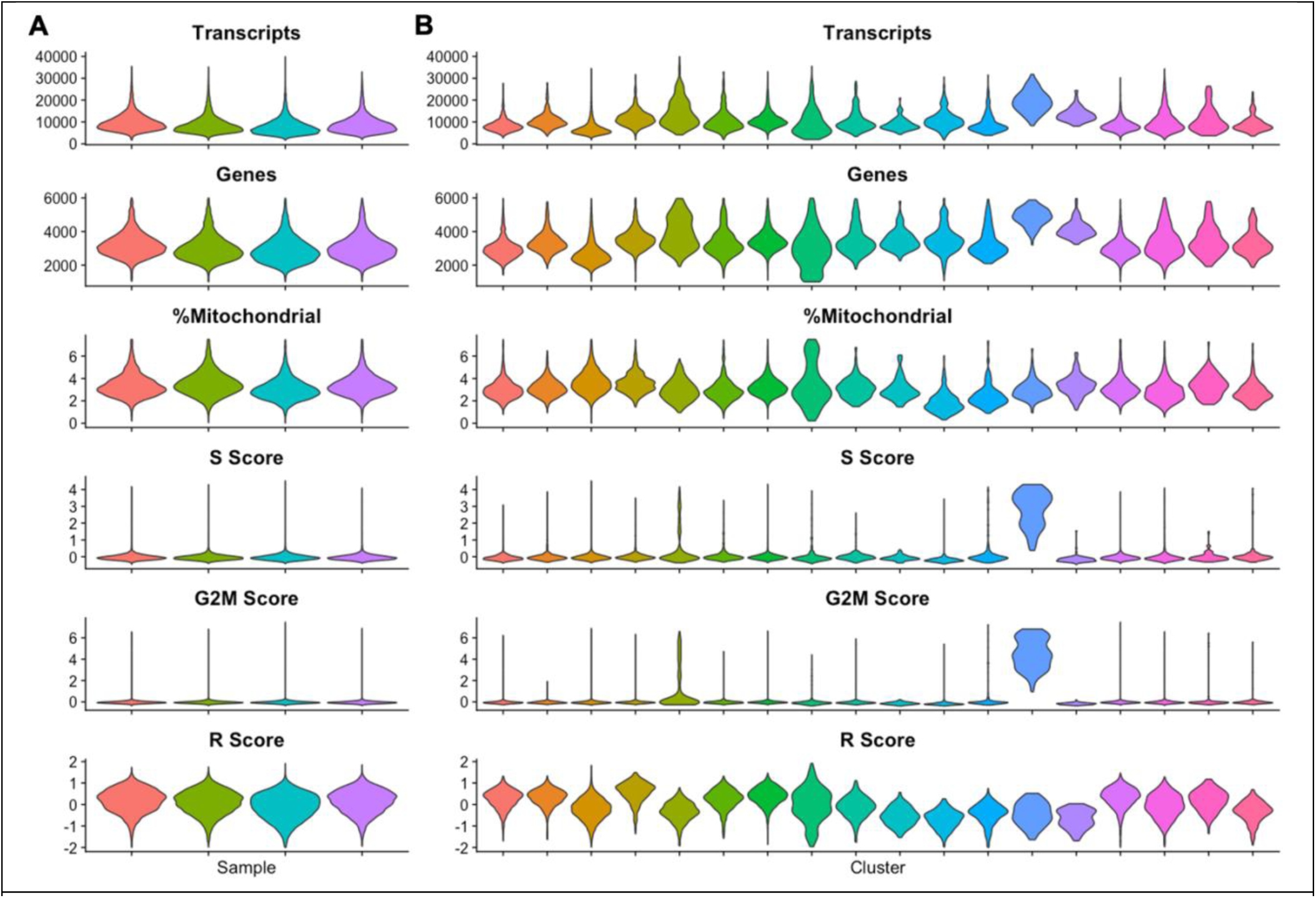
(**A**) Violin plot showing quality control metrics (from top to bottom: transcripts per cell, genes per cell, percent mitochondrial reads per cell, S Score, G2M Score, and R Score) for each sample (x-axis) processed on the 10x Genomics Chromium instrument. (**B**) Violin plot showing quality control metrics for each cell type (x-axis).

**Supplementary Figure 4.**
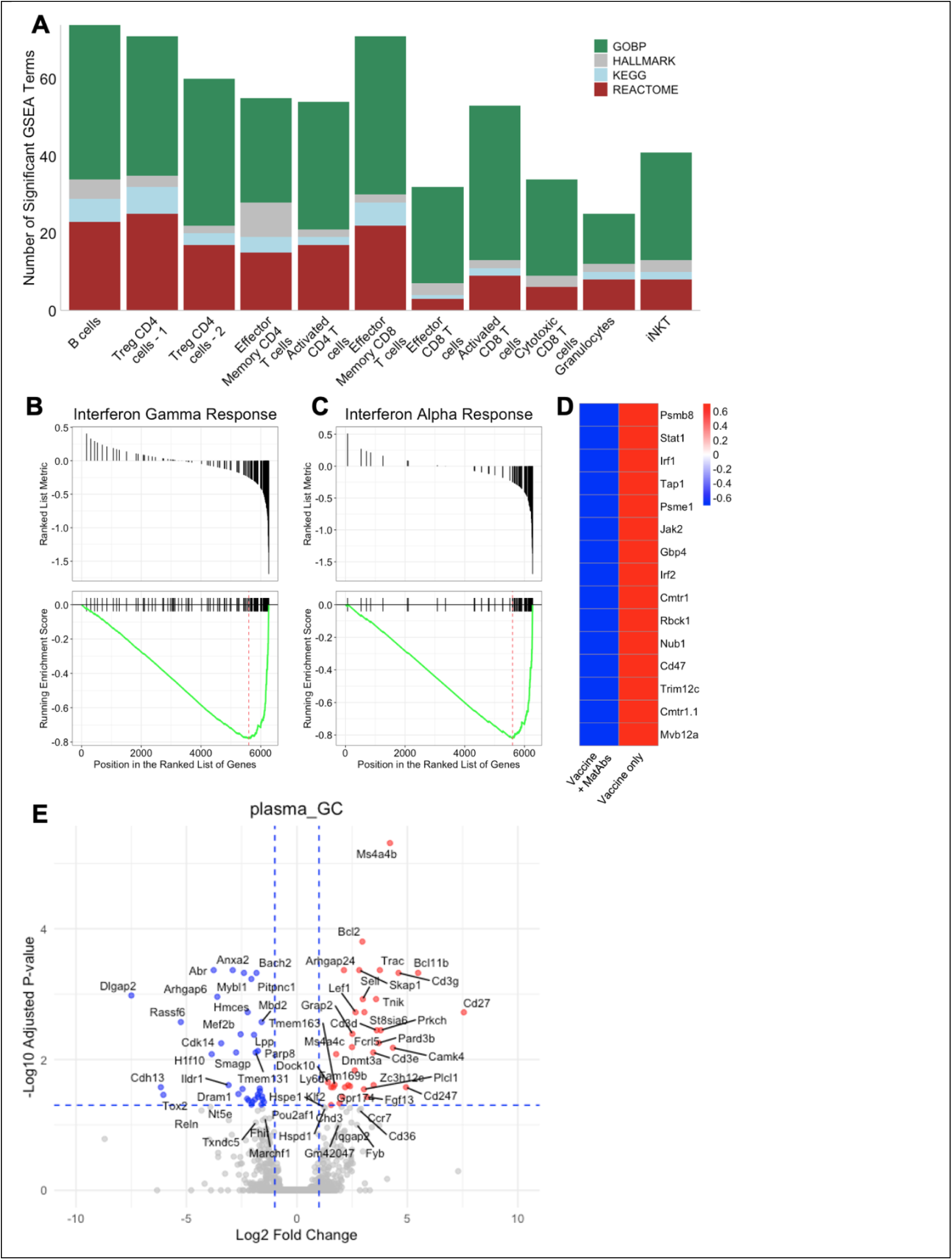
(**A**) Counts of the strongest significantly enriched gene sets with an absolute normalized enrichment score (NES) >2 across the largest cell clusters. (**B**) GSEA plot of hallmark interferon gamma response in activated CD4 T cells. (**C**) GSEA plot of hallmark interferon alpha response in activated CD4 T cells. (**D**) Heatmap of top 10 core enrichment genes taken from the gene set Hallmark Interferon Gamma and Hallmark Interferon Alpha Response between vaccine + MatAb and vaccine only; expression values row-normalized; unique genes from both pathways plotted. (**E**) Volcano plot for differentially expressed genes in vaccine + MatAb compared to vaccine only (MatAb/naïve) in plasma and GC cells. Up (red) and downregulated (blue) genes with absolute log2 fold change >1 and adjusted p-value >0.05 shown; -log10 adjusted p-value shown.

