## Supplementary figures and images for "Mechanisms of maternal antibody interference to rotavirus vaccination"

### Figure S1

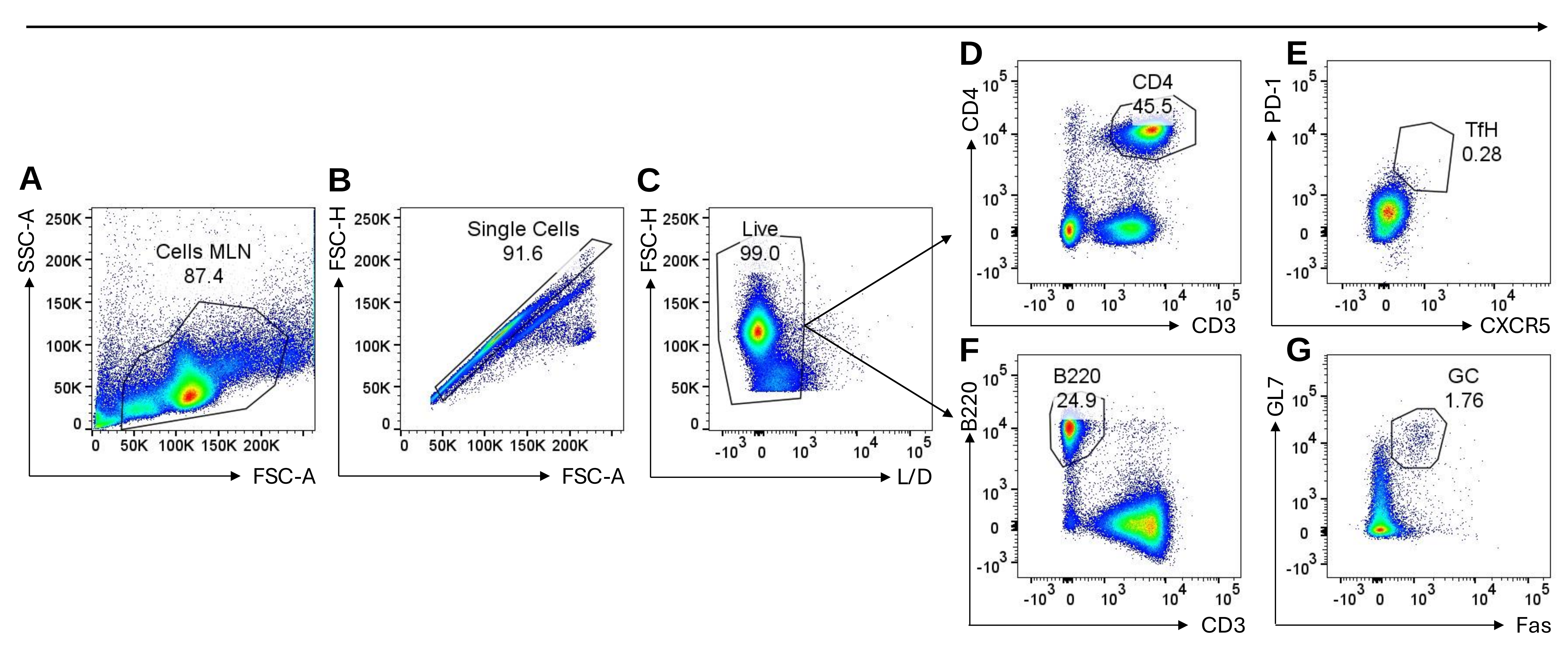

### Figure S2

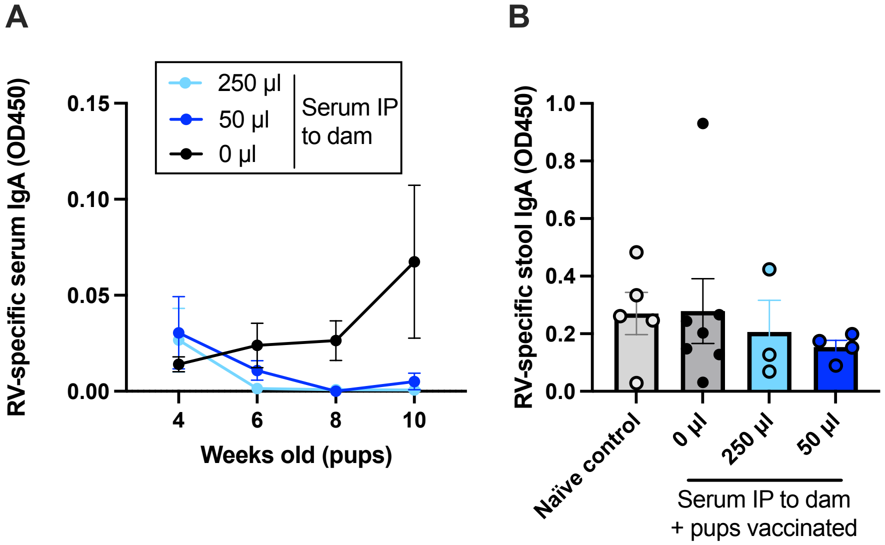

### Figure S3

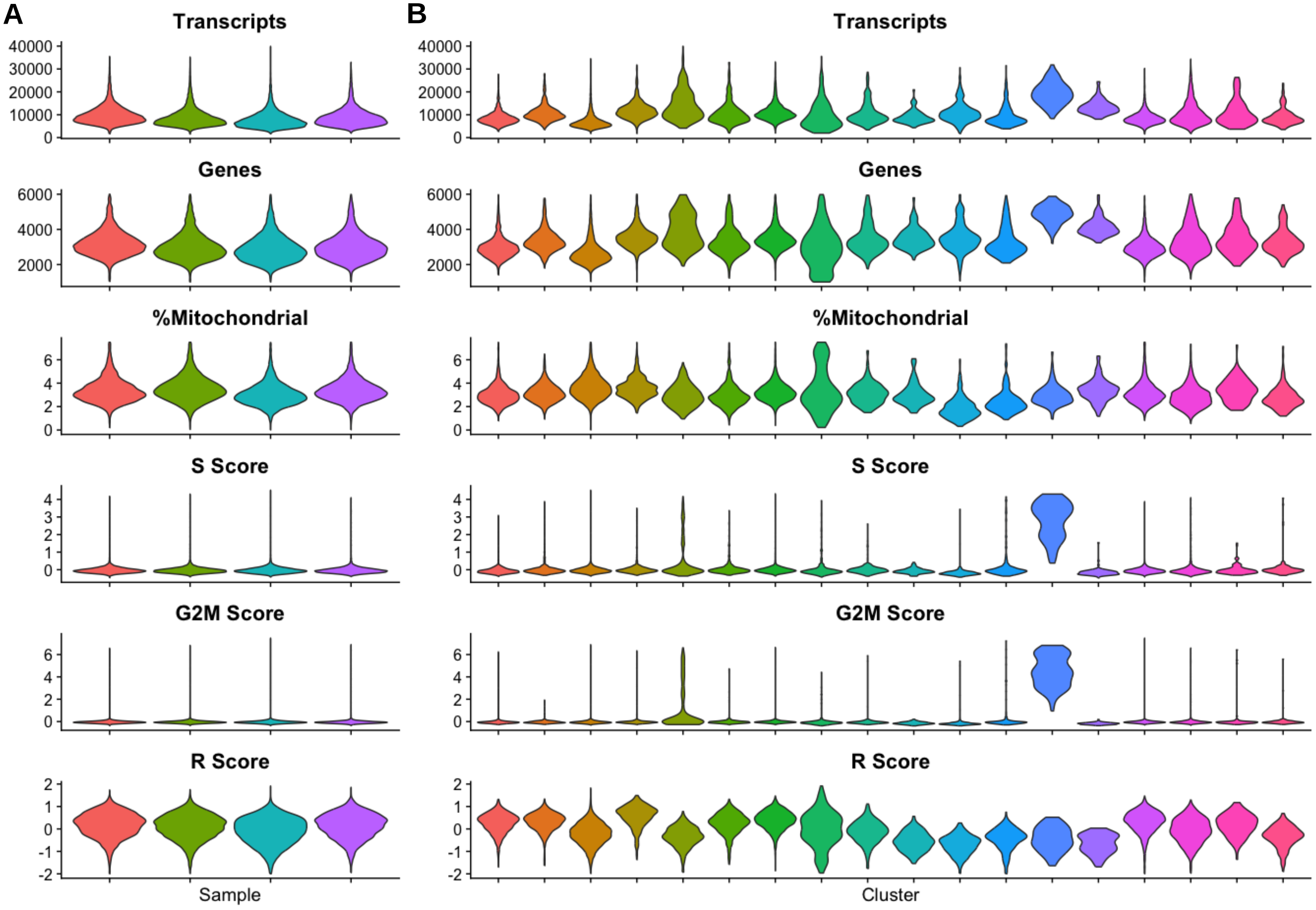

### Figure S4

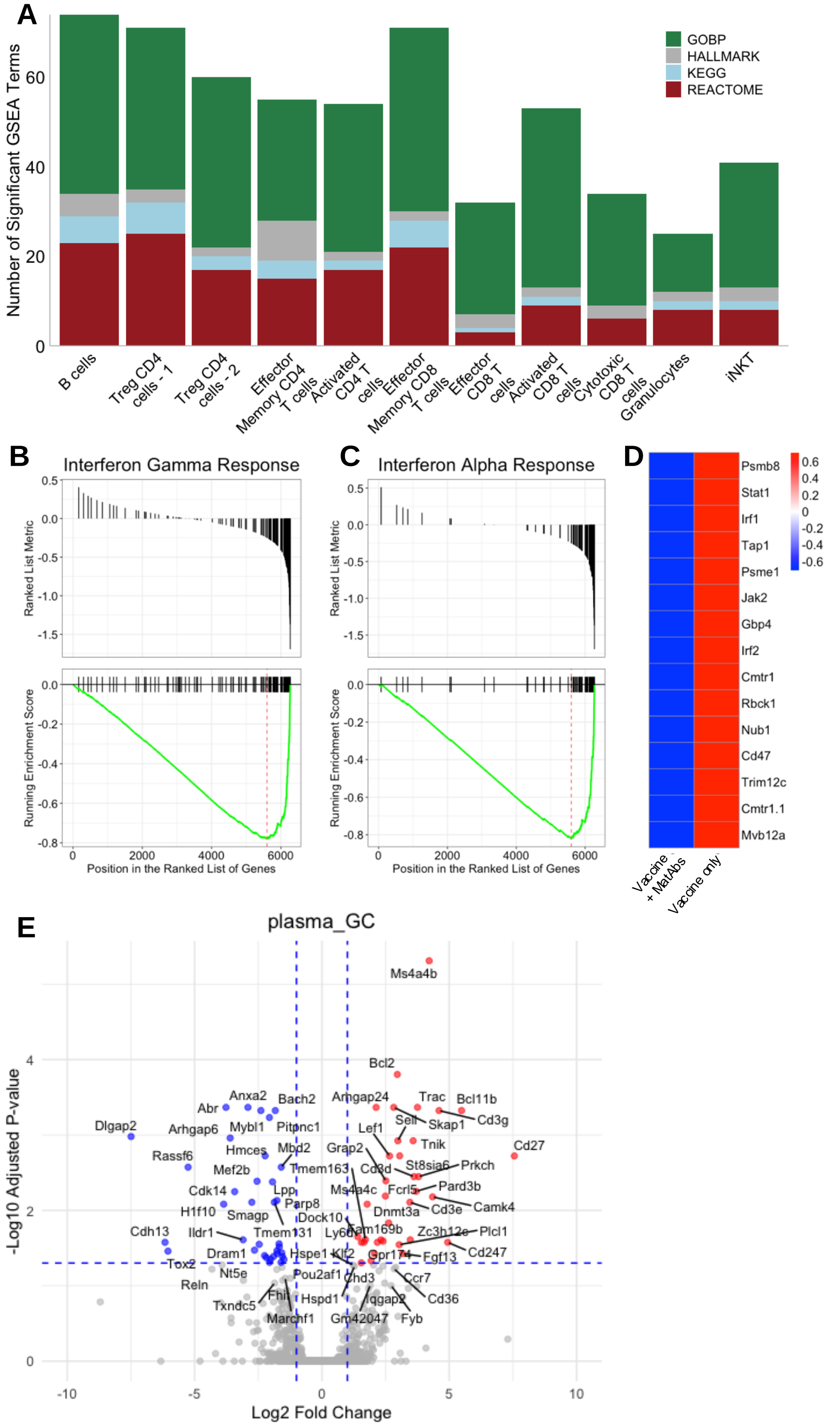
